# Large-scale discovery of recombinases for integrating DNA into the human genome

**DOI:** 10.1101/2021.11.05.467528

**Authors:** Matthew G. Durrant, Alison Fanton, Josh Tycko, Michaela Hinks, Sita S. Chandrasekaran, Nicholas T. Perry, Julia Schaepe, Peter P. Du, Peter Lotfy, Michael C. Bassik, Lacramioara Bintu, Ami S. Bhatt, Patrick D. Hsu

## Abstract

Recent microbial genome sequencing efforts have revealed a vast reservoir of mobile genetic elements containing integrases that could be useful genome engineering tools. Large serine recombinases (LSRs), such as Bxb1 and PhiC31, are bacteriophage-encoded integrases that can facilitate the insertion of phage DNA into bacterial genomes. However, only a few LSRs have been previously characterized and they have limited efficiency in human cells. Here, we developed a systematic computational discovery workflow that identifies thousands of new LSRs and their cognate DNA attachment sites by. We validate this approach via experimental characterization of LSRs in human cells, leading to three classes of LSRs distinguished from one another by their efficiency and specificity. We identify landing pad LSRs that efficiently integrate into synthetically installed attachment sites orthogonal to the human genome, human genome-targeting LSRs with computationally predictable pseudosites, and multi-targeting LSRs that can unidirectionally integrate cargos at with similar efficiency and superior specificity to commonly used transposases. LSRs from each category were functionally characterized in human cells, overall achieving up to 7-fold higher plasmid recombination than Bxb1 and genome insertion efficiencies of 40-70% with cargo sizes over 7 kb. Overall, we establish a paradigm for large-scale discovery of microbial recombinases and reconstruction of their target sites directly from microbial sequencing data. This strategy provides a rich resource of over 60 experimentally characterized LSRs that can function in human cells and thousands of additional candidates for large-payload genome editing without exposed DNA double-stranded breaks.

## INTRODUCTION

The ability to control long DNA molecules has fundamentally relied on enzymes derived from the microbe-phage interface (Faure et al. 2019; Salmond and Fineran 2015). Manipulation of eukaryotic genomes, particularly the integration of multi kilobase DNA sequences, remains challenging and limits the fast-growing fields of synthetic biology and cell engineering. Current gene integration approaches rely on DNA double-stranded breaks (DSBs) to direct cellular DNA repair pathways such as homologous recombination (HR). These approaches generally suffer from low insertion efficiency, high indel rates, and cargo size limitations. Despite important advances in optimizing HR in specific contexts (Vaidyanathan et al. 2021; De Ravin et al. 2021), the efficiency of integration typically decreases exponentially as the size of the insertion increases beyond 1 kilobase (kb) (Kung et al. 2013; Perez et al. 2005; Lee et al. 2016). Furthermore, HR-based gene editing is not feasible in post-mitotic cells and formation of DSBs is toxic in many primary cell types, sometimes leading to undesired deletions and complex rearrangements (Kosicki, Tomberg, and Bradley 2018) as well as activation of p53 (Haapaniemi et al. 2018).

Recombinases are also derived from the microbe-phage arms race and hold unique promise to address these limitations. These integrase systems have naturally evolved to catalyze DNA mobility to independently transfer genetic material from one organism to another without relying on endogenous genetic repair machinery. They are capable of catalyzing target cleavage, strand exchange, and DNA rejoining within their synaptic complex, enabling site-specific gene editing without requiring any cellular cofactors. Furthermore, because they have naturally evolved to deliver long stretches of a viral genetic code, they can exhibit large payload capacity. As a result, recombinases exhibit natural mechanistic advantages over nuclease-mediated genome editing.

Large serine recombinases (LSRs) have already yielded useful genome engineering tools, such as Bxb1 and PhiC31 (Smith 2015; Jusiak et al. 2019). Bxb1 has been used primarily as a landing pad integrase, where DNA cargos are integrated site-specifically into a pre-installed attachment site in a target genome (Inniss et al. 2017), while PhiC31 has been exploited directly for genome targeting into pseudosite loci that resemble its native attachment sites (Keravala et al. 2006; Groth et al. 2000). A major advantage of LSRs over other emerging technologies is that there is no obvious upper limit on the size of the donor DNA, with reports demonstrating successful 27 kb integration into mammalian cells (Duportet et al. 2014). While these features make LSRs highly attractive genome editing tools, the practical application of existing LSRs has been limited by several factors including low efficiency without antibiotic selection (Jusiak et al. 2019; J. Sivalingam et al. 2010).

Most efforts to improve LSR technologies have focused on enhancing the few known recombinases through processes such as directed evolution, protein fusion, domain swapping, and delivery optimization (Sclimenti, Thyagarajan, and Calos 2001; Karow et al. 2011; Farruggio and Calos 2014; Guha and Calos 2020). The advent of extensive microbial genome and microbiome sequencing efforts has enhanced the ability to discover millions of new genes (Shendure et al. 2017; Almeida et al. 2021). The abundance of both sequenced genomes and LSR proteins in nature provides a new opportunity to develop new systems that may already have evolved a desired property, as opposed to relying on laborious protein engineering.

Here, we sought to expand the LSR toolbox by computationally identifying and experimentally characterizing LSRs with innate capacity to integrate in the human genome. Reasoning that LSRs carried within mobile genetic elements (MGE) may be responsible for MGE mobility, we first searched for MGEs harboring thousands of diverse LSRs. Our group recently developed an approach that enables the prediction of boundaries of MGEs harboring LSRs in a highly precise and automated fashion, allowing systematic reconstruction of their cognate DNA recognition sites (Durrant et al. 2020). This enabled us to greatly increase the scale of our computational LSR search relative to previous methods (Yang et al. 2014).

We categorize these LSRs according to three separate technological applications: 1) landing pad LSRs that can integrate efficiently and specifically into a pre-installed attachment site (a landing pad), 2) genome-targeting LSRs with predictable specificity for endogenous sites in the human genome, and 3) multi-targeting LSRs that efficiently and unidirectionally integrate into multiple target sites with higher specificity than transposase systems. LSRs from each category were validated and functionally characterized in human cells, overall achieving up to 7-fold higher plasmid recombination than Bxb1 and genome insertion efficiencies of 40-70% with cargo sizes over 7 kb. Taken together, we establish a platform for systematic identification of integrase systems and demonstrate new LSRs as highly efficient systems for larger-scale genome editing applications.

## RESULTS

### Systematic discovery and classification of large serine recombinases and their target site specificities

LSRs canonically recombine two distinct DNA attachment sites natively found on an invading phage genome (attP) and a target bacterial genome sequence (attB). Upon phage insertion, these attachment sites are recombined into attL and attR sequences that mark the integration boundaries. We sought to systematically identify LSRs contained within MGEs and their attachment sites using a comparative genomics approach on sequence databases of clinical and environmental bacterial isolate genomes. First, we identified candidate recombinases across 194,585 bacterial isolate genomes that were contained within integrated MGE boundaries (**Fig. 1A**). We used the attL and attR sequences to reconstruct the original attP and attB attachment sites for 12,638 candidates. After applying quality control filters such as LSR coding sequence length or distance to the MGE edge, our final dataset of LSR-attachment site predictions included 6,207 unique LSRs (1,081 50% amino acid identity clusters) and cognate attachment sites. These candidates belonged to 20 host phyla, indicating good representation of published bacterial assemblies (**Fig. S1A**).

**Figure 1.**
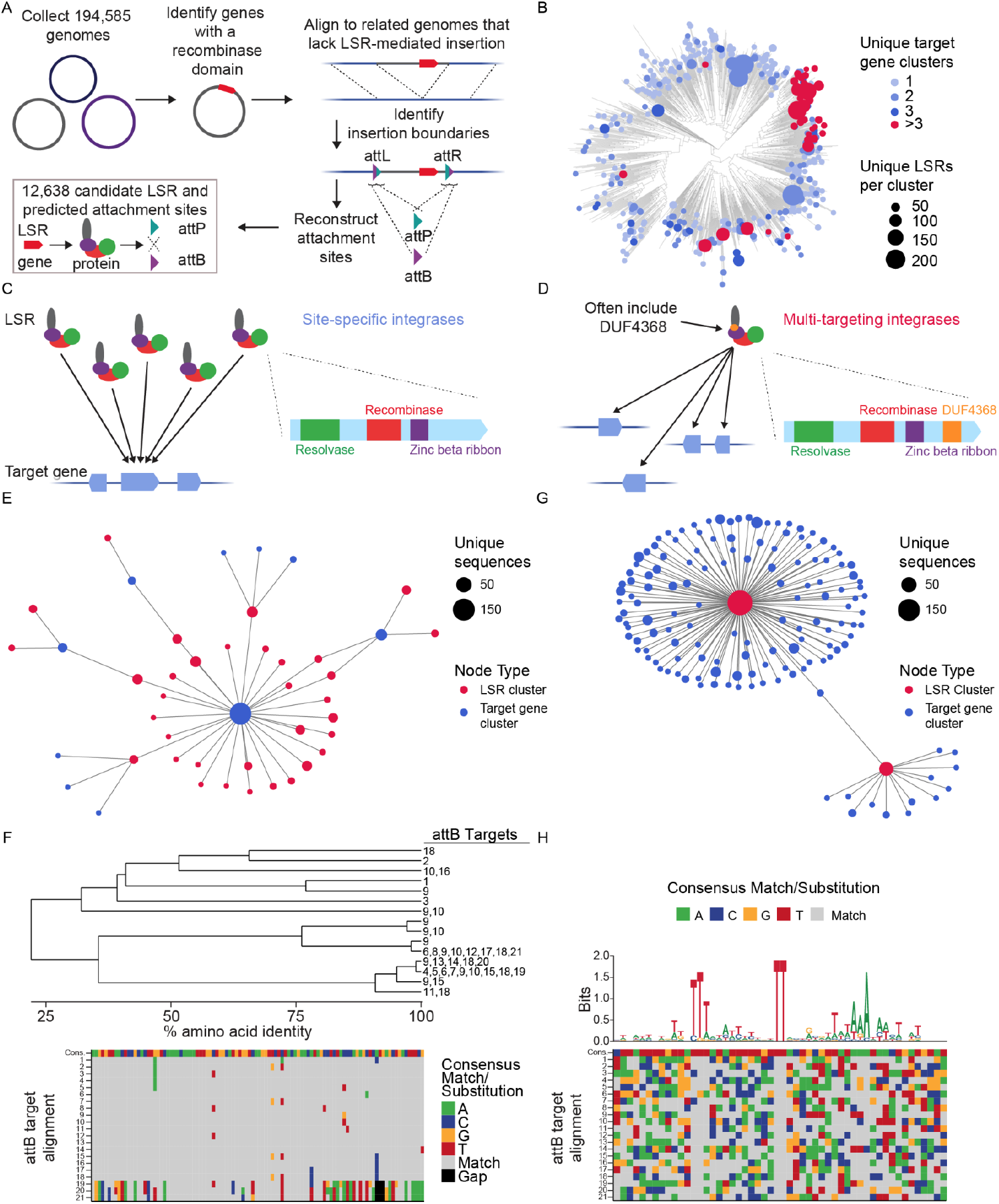
Systematic discovery and classification of large serine recombinases and their target site specificities. **A**. Schematic of computational workflow for systematic identification of LSRs and target DNA sequences, or attachment sites. Gene encoding for the recombinase domain shown as red rectangle. In the LSR protein, same notation as in panel C: recombinase domain (red), resolvase (green), Zinc betta ribbon (purple), the C-terminal (gray). **B**. Phylogenetic tree of representative orthologs across LSRs clustered at 50% identity, annotated according to predicted target specificity of each LSR family. Unique target gene clusters are the number of predicted target gene families, dots scaled to indicate the number of unique sequences found in each LSR cluster. **C**. Schematic of technique to identify site-specific LSRs that target a single gene cluster. The typical domain architecture of a site-specific LSR is illustrated. **D**. Schematic of technique to identify multi-targeting LSRs. Briefly, if a single cluster of related LSRs (clustered at 90% identity) integrate into multiple diverse target gene families (clustered at 50% identity), then the LSR cluster is considered multi-targeting. The typical domain architecture of a multi-targeting LSR is illustrated, commonly including a domain of unknown function. **E**. Example of an observed network of predicted site-specific LSRs found in our database. Each node indicates either an LSR cluster (red) or a target gene cluster (blue). Edges between nodes indicate that at least one member of the LSR cluster was found to integrate into at least one member of the target gene cluster. **F**. Example of a hierarchical tree of diverse LSR sequences that target a set of closely related attB sequences. Numbers indicate the attB sequences that are targeted by each LSR. Bottom is the alignment of related attB sequences. **G**. Example of an observed network of predicted multi-targeting LSRs. Node colors and sizes are the same as in (D). **H**. Schematic of an alignment of diverse attB sequences that are targeted by a single multi-targeting LSR. Each target sequence is aligned with respect to the core TT dinucleotide. Sequence logo above the alignment indicates conservation across target sites, a proxy for the sequence specificity of this particular LSR. The alignment is colored according to the consensus, the same as in (E).

Next, we sought to bioinformatically predict the site specificity of our candidate LSRs (**Fig. 1B**). To do this, we compared integration patterns across LSR clusters, grouping attB sites by the genes that they overlapped with. We reasoned that if many distantly-related LSRs appear to target similar integration sites, it is likely that these LSRs would be site-specific (**Fig. 1C**). We predict that 82.8%-88.3% of clusters are site-specific or have intermediate site-specificity, where the total number of unique target genes is 1, 2, or 3, depending on strictness of criteria used. Conversely, if we saw LSR clusters that targeted many distinct integration sites, we then classified them as “multi-targeting”, meaning that they either had relaxed sequence specificity and/or they evolved to target sequences that occurred at multiple different sites in their host organisms (**Fig. 1D**). One significant clade of multi-targeting LSRs are predicted to integrate into more than 3 target gene clusters, suggesting that this was an evolved strategy inherited from a single ancestor (**Fig. 1B, S1A**). We find that most (63%) LSRs in this clade contain DUF4368, a Pfam domain of unknown function, which is rarely (0.73%) found in site-specific LSRs (**Fig. S1A**), and that the clade includes previously described LSRs in the TndX-like transposase subfamily (H. Wang and Mullany 2000; Adams et al. 2004; Wang Hongmei, Smith Margaret C. M., and Mullany Peter 2006).

We found many examples of distantly related LSRs that targeted the same gene clusters, including a network of diverse LSR clusters that primarily target a single gene cluster (**Fig. 1E**,**F**). Homologs of this particular gene, annotated as an ATP-dependent protease, are one of the most commonly targeted genes, being targeted by 12.4% of all predicted site-specific integrases (**Fig. S1B**). In another striking example, we found a diverse set of 33 unique LSRs (15 99% amino acid identity clusters, 6 50% identity clusters) that all target a single conserved site, the CDS sequence of a Prolyl isomerase (**Fig. 1F**). Upon aligning the LSR candidates that targeted this site, we found that the DNA-binding Resolvase, Recombinase, and Zn_ribbon_recom domains were more conserved than the C-terminus, which primarily plays a role in protein-protein interactions (McEwan, Rowley, and Smith 2009), suggesting that conservation in DNA-binding domains among these LSRs may reflect their shared target site specificity (**Fig. S1C**).

We also identified clusters of LSRs where highly related orthologs integrate into divergent targets (**Fig. 1G**). Several of these multi-targeting LSRs have large numbers of associated attB target sites, which allows us to infer their sequence specificity computationally from our database. In one example, we found a single multi-targeting integrase that targets 21 distinct DNA sites. An alignment of these target sites revealed a conserved TT dinucleotide core with 5′ and 3′ ends enriched for T and A nucleotides, suggesting that this particular LSR most likely has relaxed sequence specificity overall (**Fig. 1H**). Other multi-targeting LSRs appear to have distinct target site motifs, including several with more complex motifs than short, AT-rich sequences (**Fig. S1D**). Overall, these analyses demonstrated the power of large-scale recovery of LSRs and attachment sites, as they provide insight into the differences in targeting specificity across the diversity of serine integrases. Further, they suggest that we may be able to predict if these LSRs would target, or avoid off-targets, in the human genome if applied as genome engineering tools.

### Efficient landing pad LSRs in human cells

One valuable application for LSRs is the specific delivery of genetic cargo to an introduced landing pad site. To validate our computational predictions, we synthesized human-codon optimized LSRs and their predicted attP and attB sequences, and validated recombination activity in human cells via a plasmid recombination assay (**Fig. 2A**). In this assay, the attP plasmid contains a promoterless mCherry, which gains a promoter upon recombination with the attB plasmid. Out of 17 candidates, we identified 15 functional candidates (88%), defined as having greater mCherry MFI values than their attP-only controls (**Fig. 2B, C**). 13 candidates had favorable recombination efficiency relative to PhiC31 while 3 were superior to Bxb1 (**Fig. 2B**). These mCherry MFI results were in general agreement with the percentage of mCherry^+^ cells, a metric unaffected by differences in transcription, delivery, and frequency of recombination events across cells (**Fig. S2A**). We tested activity of a subset of LSRs with different attachment site combinations and found that they are highly orthogonal and require their cognate attachment sites (**Fig. 2D**). These results indicate that our LSR discovery approach yields a high validation rate.

**Figure 2.**
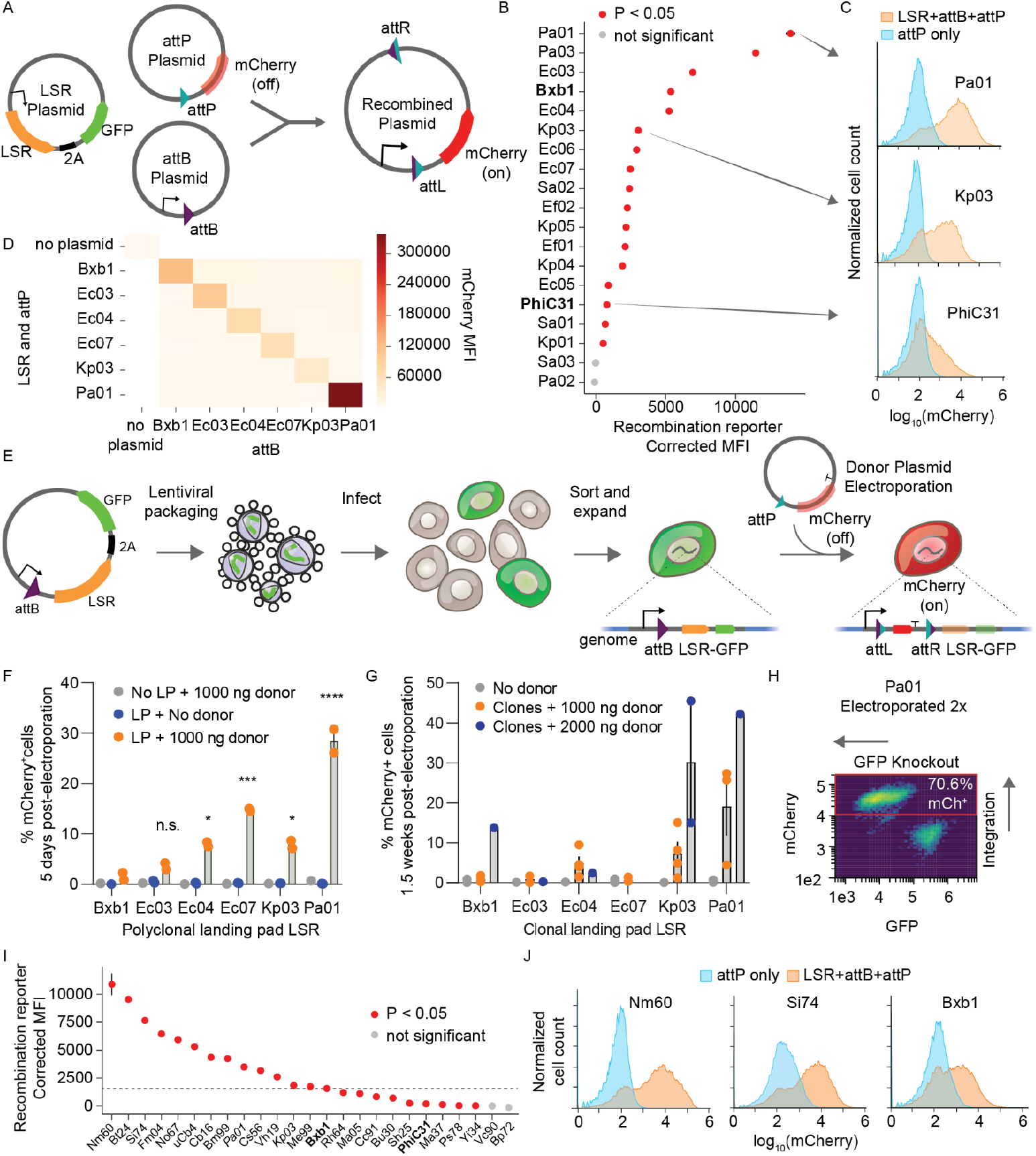
Development of efficient and specific recombinases for human landing pads. **A**. Schematic of plasmid recombination assay. Cells are co-transfected with LSR-2A-EGFP, promoter-less attP-mCherry, and pEF-1α-attB. Upon recombination, mCherry gains the EF-1α promoter and is expressed. **B**. Plasmid recombination assay of predicted LSRs and att sites in HEK293FT cells, shown as mCherry mean fluorescence intensity (MFI) corrected by subtracting the average MFI of attP-only negative controls. Data are mean (n=3) ± s.d. P-value determined by one-tailed t-test. **C**. Example mCherry distributions for all three plasmids (LSR+attB+attP) compared to the attP-only negative control. **D**. Plasmid recombination assay between all pairs of LSR+attP and attB in K562 cells (n=1 electroporation replicate). **E**. Schematic of genomic landing pad assay. An EF-1α promoter, attB, and LSR are integrated into the genome of K562 cells via lentivirus. Cells are then electroporated with the attP-mCherry donor plasmid. Upon successful integration into the landing pad, mCherry is expressed, and the LSR and GFP are knocked out. **F**. Efficiency of promoterless-mCherry donor integration into a polyclonal genomic landing pad (LP) K562 cell line, measured after 5 days (n=2 independently transduced and then electroporated biological replicates). Asterisks show statistical significance for landing pad plus donor conditions compared to Bxb1 (one-way ANOVA with Dunnett’s multiple comparisons test, * is P < 0.05, *** is P < 0.001, **** is P < 0.0001, n.s. is not significant). **G**. Donor plasmid integration into clonal landing pad cell lines electroporated with 1000 ng donor plasmid (10 days after electroporation) or 2000 ng donor plasmid (11 days after electroporation). Each point is a different clonal K562 cell line carrying the landing pad and LSR corresponding with the donor, error = SEM. 1000 ng Pa01 is significantly more efficient than 1000 ng Bxb1 comparing between conditions with donor electroporation (P < 0.005, one-way ANOVA, n=3 clonal cell lines for Pa01 and n=4 clonal cell lines for others at 1000 ng dose, n=2 clonal cell lines for Kp03 and n=1 clonal cell line for others at 2000 ng dose). **H**. Flow cytometry showing knockout of LSR-GFP and integration of mCherry in the same cells. Pa01 clonal landing pad line mixed with 1000 ng donor was electroporated twice in rapid succession to increase delivery, resulting in >70% mCherry^+^ cells after 11 days. **I**. Plasmid recombination assay for a batch of LSRs selected for higher quality (see Methods) in HEK293FT cells. Data are mean (n=3 transfection replicates) ± s.d. Controls are labeled in bold and LSRs from the previous batch are labeled in italics.P-value determined by one-tailed t-test. **J**. Representative mCherry distributions for all three plasmids (LSR+attB+attP) compared to the attP-only negative control.

To test the efficiency of integration into the endogenous human genome, we next generated cell lines with genomic knock-in of attB-containing landing pads. We developed a promoter trapping approach where successful recombination would lead to mCherry expression and loss of LSR and GFP expression (**Fig. 2E**). All of the tested LSRs could integrate with measurable efficiency, four of which were significantly more efficient than Bxb1 (**Fig. 2F**). Next, we assessed the stability of these polyclonal landing pad cell lines. While GFP expression in landing pads such as Pa01 remain constant over time, others lose GFP expression, suggesting the landing pad can be transcriptionally silenced or genetically unstable in a manner that depends on the attB or LSR (**Fig. S2B**). Overall, these results confirmed that new LSRs can efficiently integrate donor cargo into human chromosomal DNA at landing pads.

Landing pad applications may necessitate a single genomic integration site in all cells. To develop single position landing pad lines, we integrated the landing pad LSR-GFP construct via low MOI lentiviral infection, clonally expanded GFP^+^ cells, and electroporated with an attP-mCherry donor plasmid. We tested four integrase candidates and found that Pa01 performed better than Bxb1 across multiple independent clones in terms of the percentage of cells that were stably mCherry positive after 1.5 weeks (average of 19% versus 1%) (**Fig. 2G**). With a doubled donor DNA dose, Pa01 reached 42% efficiency (**Fig. 2G**). Electroporating cells with donor plasmid twice in rapid succession increased integration efficiency to over 70%, suggesting efficiency was primarily limited by donor delivery (**Fig. 2H**). These cells were also GFP-negative, consistent with the desired outcome of an mCherry donor being integrated into the landing pad and knocking out the LSR-GFP cassette in the process.

Reasoning that shorter attachment sites would facilitate the installation of landing pads, we next set out to identify the minimal attB sequences necessary for recombination. Previous characterization of the Bxb1 attB identified a sequence as short as 38 bp as being necessary for integration, but our computational pipeline conservatively predicts 100 bp attB sequences initially. We determined a minimum 33 base pair attB for efficient Pa01 recombination (**Fig. S2C)** and observed efficient recombination for Kp03 down to a 25 base pair attB (**Fig. S2D**). At such short lengths, these attachment sites can easily be installed during cloning and cell engineering methods while minimizing perturbation of endogenous sequence.

Efficient landing pad recombinases could be especially useful for multiplex gene integration. This could be achieved by using several of the new LSRs in parallel, given their demonstrated orthogonality (**Fig. 2D**). Bxb1 has been shown to contain a modular dinucleotide core in its attachment sites that enable orthogonal integrations such that the same LSR can direct multiple cargoes to landing pads that differ only by their core dinucleotides (Ghosh, Kim, and Hatfull 2003). We tested the ability to substitute core dinucleotides using our plasmid recombination assay for one of our candidates, Kp03, confirming the orthogonality conferred by altering the central dinucleotides (**Fig. S2E**).

We then investigated the specificity of these LSRs by transfecting LSRs and mCherry donors and measuring mCherry expression over time, as episomal donor plasmid dilutes and stable integration remains. At day 18, Pa01 showed no evidence of integration above background, while Kp03 did have elevated percentages of mCherry^+^ cells, suggesting that it has off-target pseudosites (**Fig. S2F**). To identify these sites, we modified an NGS assay originally developed for analyzing Cas9 edits and translocations for use as an LSR integration site mapping assay (**Fig. S2G, H**) (Giannoukos et al. 2018; Danner, n.d.). First, we quantified the percentage of off-target integrations relative to on-target integrations in landing pad cell lines (**Fig. S2I**). This assay detected off-target integrations for all LSRs, including Bxb1 (3.48%), Pa01 (0.47%), and Kp03 (15.5%). Additionally, we developed target site sequence motifs from precise integration sites (**Fig. S2J**). These motifs validate the experimentally determined minimal attachment site length and demonstrate the highly conserved dinucleotide core. Together, these results establish Pa01 as a more efficient and comparably specific landing pad LSR in comparison to Bxb1, and provide an expanded set of additional landing pad LSRs.

Finally, we selected a second batch of 21 LSRs from our database, prioritizing those with low BLAST similarity between their attB/P sites and the human genome, and applying stringent quality thresholds (see Methods). We found 17 out of 21 (81%) of them were functional in the plasmid recombination assay, providing further validation of the computational pipeline for identifying functional candidates. Promisingly, 16 candidates were more efficient than PhiC31 while 11 were superior to Bxb1 (**Fig. 2I, J**). Our fluorescence integration assay in wild-type cells identified 4 LSRs with minimal off-target integrations, nominating Si74 and Nm60 as two top LSRs with high recombination efficiency and genomic specificity, two of the most critical features for therapeutic applications of these integrase technologies (**Fig. S2K**).

### Landing pad LSRs enable parallel reporter assay

To explore the utility of landing pad LSRs in functional genomics, we established a parallel reporter assay (PRA) that tests the capacity of synthetic enhancers to activate a transcriptional reporter integrated in the genome (**Fig. 3A**). PRAs have recently become an effective means of studying diverse molecular elements including enhancers, promoters, and untranslated regions (Haberle et al. 2019; King et al. 2020; Rabani et al. 2017). However, because PRAs can be adversely affected by genomic position effects or other forms of heterogeneity in delivery (Klein et al. 2020), LSR landing pads can enable efficient and homogenous stable integration of PRA reporters into a single genomic position (Maricque, Chaudhari, and Cohen 2018; Cao et al. 2021; Matreyek et al. 2020).

**Figure 3:**
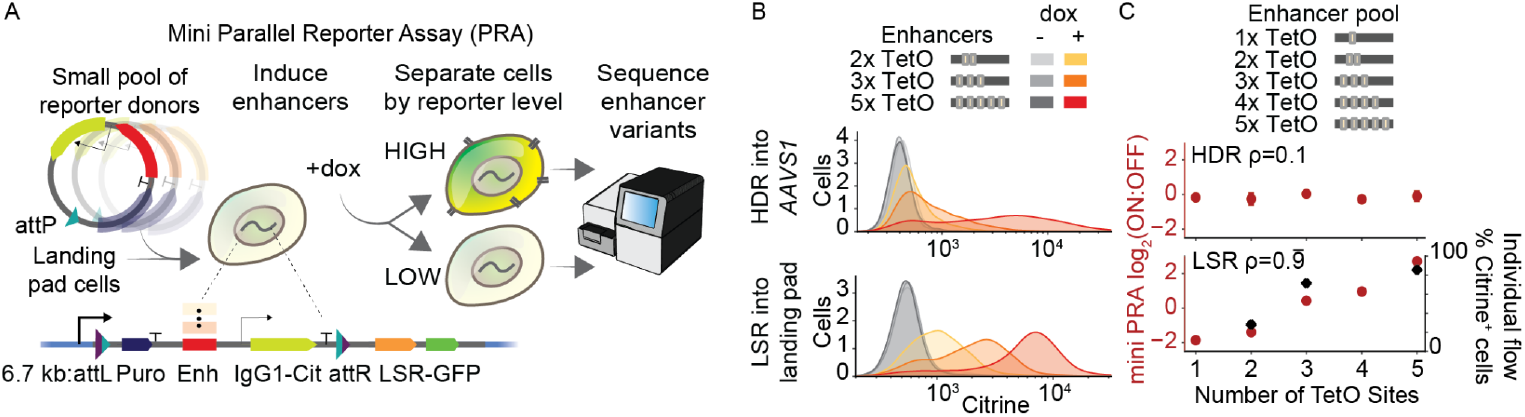
Parallel reporter assays with landing pad recombinases. **A**. Schematic of mini parallel reporter assay (PRA) using the landing pad. A pool of reporter donor plasmids with varied synthetic enhancers containing TetO sites is integrated into the landing pad by the LSR, selected for using puromycin, and then reporter activation is induced using doxycycline which causes rTetR-VP48 to bind TetO. Highly- and lowly-activated cells are magnetically separated, the enhancers are sequenced from genomic DNA in each cell population, and a ratio of sequencing reads is computed as a measurement of enhancer strength. **B**. Individual synthetic enhancer reporters with a varied number of TetO transcription factor binding sites were each cloned with homology arms and integrated into the AAVS1 safe harbor by HDR or cloned with attP and individually integrated into the landing pad using the Kp03 LSR. Flow cytometry measurements of the activated citrine reporter 2 days after induction with doxycycline. Due to varied voltage settings on the cytometer, the x-axes are not comparable in absolute terms (n=1 cell line replicate, and second replicate is shown in **Fig. S3A**). **C**. A small pooled library of synthetic enhancer reporters were integrated into the AAVS1 safe harbor by HDR or a clonal landing pad by the Kp03 LSR and measured as a PRA by separation and sequencing (n=2 integration replicates for HDR, n=1 integration replicate for LSR). ρ is the Spearman correlation between the PRA measurement of enhancer strength and the number of TetO sites in the enhancer. For the LSR, pooled measurements (left y-axis, red circles) correlate with the percentage of Citrine^+^ cells from individual reporter assays (right y-axis, black diamonds, Pearson’s *r*=0.94).

First, we individually integrated enhancer reporters containing a varied number of transcription factor binding sites into a Kp03 landing pad clonal line. As a control, we also used homology-directed repair (HDR) to integrate matched reporters into the AAVS1 safe harbor in cells lacking the landing pad. In both cases, the reporters contain a puromycin resistance gene that traps a promoter (the landing pad EF-1α promoter or the endogenous AAVS1 promoter, respectively), which we used to select for on-target integrations. As expected, we found that both HDR- and LSR-integrated reporters activated to degrees corresponding with the number of transcription factor binding sites in the enhancers (**Fig. 3B**).

Then, to test reporters in a parallelized fashion, we integrated a pooled library and performed a PRA (**Fig. 3A, C and S3A-C**). For the HDR-installed libraries, we did not see the expected positive correlation between enhancer activation strength and number of transcription factor binding sites (ρ=0.1), which could be due to integration of multiple library members at more than one AAVS1 allele per cell (**Fig. 3C**) (Weingarten-Gabbay et al. 2019). Meanwhile, for the LSR-installed libraries, we saw the expected correlations between the enhancer activation strength and number of transcription factor binding sites (ρ = 0.99), and also between the PRA and individual reporter measurements by flow cytometry (*r* = 0.94, **Fig. 3C**). We cannot rule out the possibility that the HDR-based strategy could be optimized to yield similar results. Taken together, these PRA results demonstrated that landing pads can be useful for making parallelized quantitative measurements of a library of reporters and indicated that these new landing pad LSRs could enable diverse functional genomics research applications.

### Genome-targeting LSRs can integrate into the human genome at predicted target sites

Ideally, LSRs would efficiently integrate directly into a single or limited number of target sites or pseudosites already existent at safe locations in the human genome. Historically, the integration sites of pseudosite-rich LSRs such as PhiC31 had to be experimentally discovered by transfecting the LSR into human cells and searching for the integration sites. Given the expanded size of our LSR database with defined attB and attP sequences, we reasoned we could first computationally search for LSRs that naturally target an attachment site highly similar to a single sequence in the human genome.

We used BLAST to search all attB/P sequences against the GRCh38 human genome assembly (**Fig. 4A**) and identified 856 LSRs with a highly significant match for at least one site in the human genome (BLAST E-value < 1e-3, **Fig. 4B, C**). We synthesized 103 LSRs prioritized by high BLAST match quality in the plasmid recombination assay, rather than our quality control metrics, and confirmed that 25 candidates recombined at predicted attachment sites (one-tailed t-test, P < 0.05; **Fig. S4A**). We found that 21 out of 37 (56.75%) high-quality candidates recombined as predicted, in contrast to 4 out of 64 (6.25%) low-quality candidates. We named the attP and attB sites according to their BLAST hits, with the attachment site that matched the human genome being renamed to attA (acceptor), and the other being renamed to attD (donor). The predicted target site in the human genome was renamed attH (human) (**Fig. 4A, C**), and we confirmed that several of our candidates recombined with their predicted attH sequence in the plasmid recombination assay (**Fig. S4B**).

**Figure 4.**
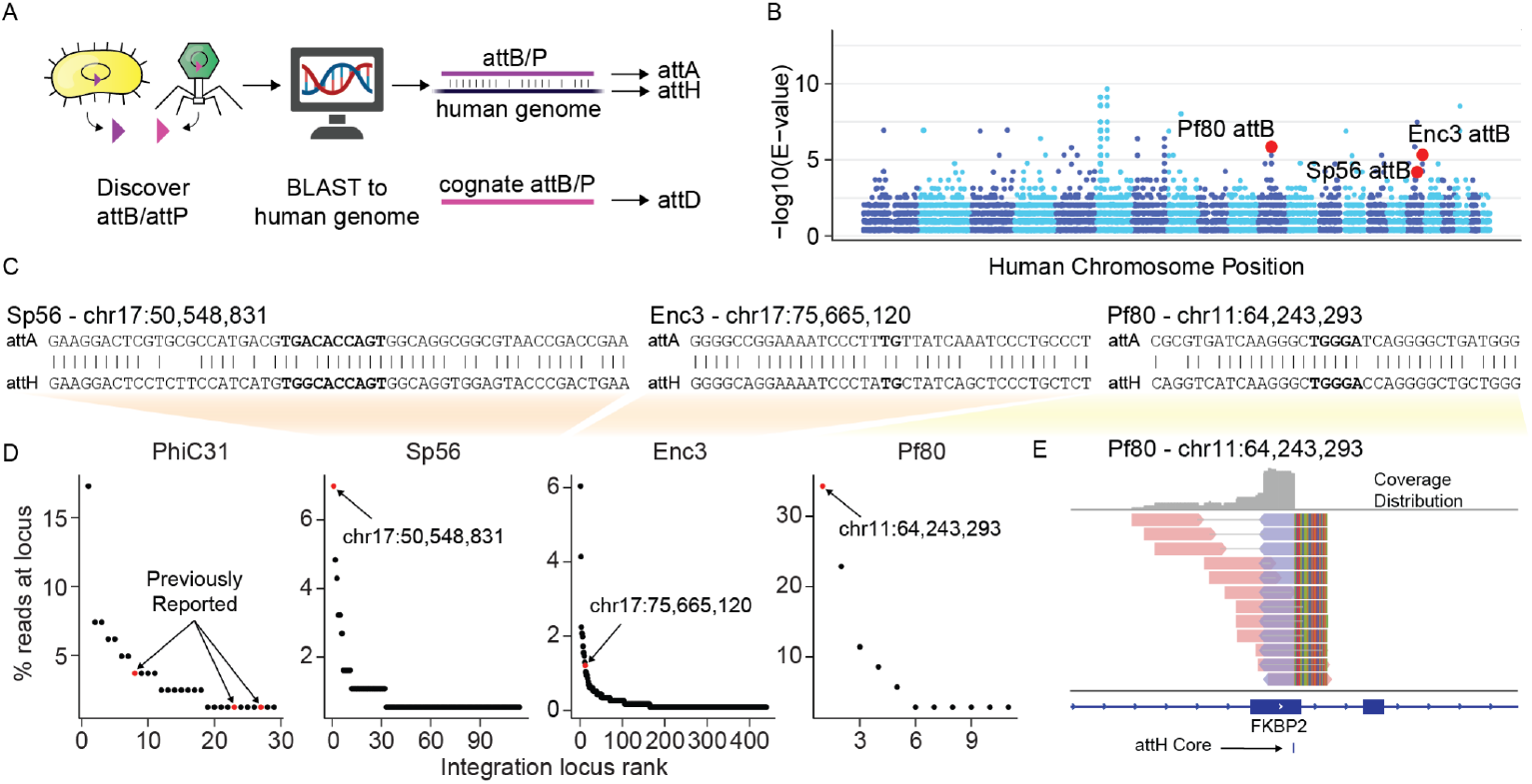
Validation of target site predictions for human genome-targeting recombinases. **A**. Schematic representation of computational strategy to identify candidates with innate affinity for the human genome. Briefly, attB and attP candidates in our database were searched against the human genome using BLAST. The attachment site with the best match to the human genome is denoted attA (acceptor), and the corresponding human genome target site is denoted attH (human). The paired attachment site is denoted attD (donor). **B**. BLAST hits of attB and attP sites that are homologous to sequences in the human genome, showing all hits that meet E < 0.01. The 22 autosomal chromosomes, starting with chromosome 1 in dark blue on the left, are shown in alternating colors with light blue every other chromosome. **C**. Schematic of BLAST alignments of the microbial attachment sites (attA) to the predicted human attachment sites (attH) for three candidates. The attachment site center is bolded, representing the portion of the native attP and attB that is identical and presumed to contain the dinucleotide core. **D**. Detected integration sites, ranked according to the number of unique reads found at each site. **E**. Reads that align to the integration sites for Pf80 in the human genome, showing the predicted attH target site. Reads that align in the forward direction are shown in red and those aligning in the reverse direction are shown in blue, with a gray line connecting paired reads.

Next, we mapped their integration sites in the human genome to test our computational predictions. For Sp56 and Pf80, the predicted target loci by BLAST were indeed the top integration sites with the most uniquely mapped reads (**Fig. 4D, E and S4C**). For Enc3, the predicted target site was among the top integration loci, although it was not the most frequently targeted locus (**Fig. 4D and S4D**). Of the tested LSRs, Pf80 had the highest predicted specificity, with 34.3% of unique reads mapping to the predicted target site (**Fig. 4D, E**). To further understand pseudosite preference, we mapped the integration sites of some genome-targeting LSRs and uncovered evidence of target site motifs (**Fig. S4E, F, G**), suggesting that these LSRs selectively recognize specific nucleotide positions within their attachment sites. Genome-targeting candidates had varying levels of efficiency, with Enc3 in particular having significantly higher efficiency than PhiC31 at 6% and <1%, respectively (**Fig. S4H**). Taken together, our computational pipeline is able to nominate serine integrases that are likely to target the human genome and predict their target site preference.

### Multi-targeting LSRs are highly efficient and unidirectional in human cells

Some serine recombinases have evolved transposition or multi-targeting capabilities, allowing them to target many different attB sites in a given prokaryotic genome (Smith and Thorpe 2002). Efficient insertion into a defined series of pseudosites would be very useful relative to semi-random integrases such as Piggybac or Sleeping Beauty transposase. We noticed that LSRs in the multi-targeting clade (**Fig. 1D, G**) often contained a domain of unknown function (DUF4368), and tested an LSR ortholog from *Clostridium perfringens* that we named Cp36. Cp36 successfully integrated an mCherry donor cargo into the genome of K562 and HEK293FT cells at up to 40% efficiency without pre-installation of a landing pad or antibiotic selection (**Fig. 5A, S5A**).

**Figure 5.**
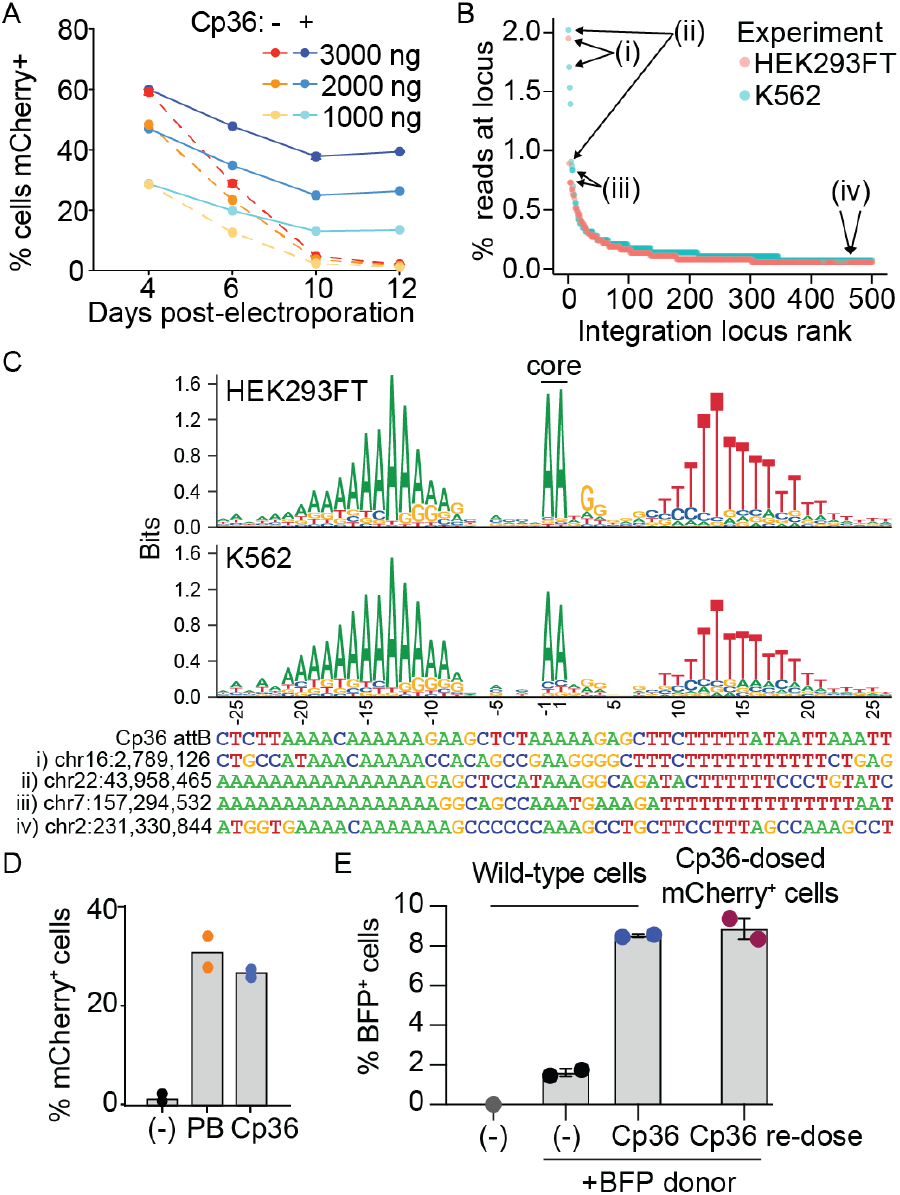
Development of highly efficient and unidirectional recombinases with defined target site motifs. **A**. Co-electroporation of LSR Cp36 and attD-mCherry donor plasmid to K562 cells. Bxb1 paired with a Cp36 attD donor was used as a negative control. The dose in ng refers to the LSR plasmid and the attD donor plasmid was delivered at a 1:1 molar ratio. Stable fluorescence in cells treated with Cp36 was measured out to 12 days post-electroporation by flow cytometry. Data are mean (n=2 transfection replicates) ± s.d. **B**. Integration site mapping assay results for Cp36 in two cell types. We define an integration locus as a sequenced integration of a donor cargo at a specific site, where sites that are within 500 bp of each other are combined into a single locus. Showing the top 500 loci across two experiments, one performed in HEK293FT cells and another performed in K562 cells. Counting uniquely mapped reads results in conservative count estimates for loci with higher coverage. The sequences of sites indicated by arrows are shown at the bottom of (C). **C**. Cp36 target site motifs and example target sequences. Precise integration sites and orientations were inferred at all loci, and nucleotide composition was calculated for the top 200 sites in the HEK293FT and K562 experiments. The core dinucleotide is found at the center. Example integration sites specified in (B) are shown below, colored according to corresponding nucleotides. **D**. Efficiency of Cp36 and Super PiggyBac (PB) for stable delivery of mCherry donor plasmid in K562 cells. The 7.2 kb donor plasmid contains the Cp36 attD and the PiggyBac ITRs. Ec03 LSR is used as a negative control that lacks an attachment site on this donor plasmid. Bars show the mean and dots show replicates (n=2 electroporation replicates). **E**. Reusability of Cp36 for serial delivery of a second fluorescent reporter (mTagBFP2). Bars show the mean, dots show replicates, error = SEM (n=2 electroporation replicates).

Using the integration site mapping assay, we identified over 2000 unique integration sites for Cp36 with the top ten loci accounting for 8.27% and 11.4% of uniquely mapped reads in HEK293FT cells and K562 cells, respectively. (**Fig. 5B, S5B**). Across these two cell types, we observed high correlation between the top integration sites (Pearson’s *r* = 0.45, P = 0.0002, **Fig. S5C**). Next, we constructed a sequence motif for Cp36 targets, which is composed of an A-rich 5′ region, followed by the AA dinucleotide core, followed by a 3′ T-rich region (**Fig. 5C**). The motif also corresponds well with our own motif prediction using only LSRs in the same 50% amino acid identity cluster as Cp36 and their cognate attB sites in our database (**Fig. S5D**).

We next compared Cp36 to the Super PiggyBac transposase, a common tool for delivering DNA cargos semi-randomly into TTAA tetranucleotides found in a target genome. We designed a 7.2 kb plasmid construct that included a Cp36 attD (donor attachment site), PiggyBac ITR sequences, and an mCherry reporter to directly compare the efficiencies of these two enzymes (**Fig. S5E**). We find that Cp36 performs at similar efficiencies to PiggyBac (26.6% and 30.9% of cells with stable integration, respectively) (**Fig. 5D**), despite Cp36 being the unaltered microbial protein sequence and Super PiggyBac being an engineered, hyperactive version of the transposase intended for genome engineering (Yusa et al. 2011).

PiggyBac is a bidirectional integrase and excisionase, resulting in both excision and local hopping of cargo upon redosing cells (W. Wang et al. 2008). It would be useful to have a single integrase that can be used repeatedly to engineer cells with multiple cargos. To test if Cp36 could be re-used to integrate a second gene, we generated mCherry^+^ cells via Cp36 and puromycin selection, then re-electroporated them with Cp36 and a BFP donor. After 13 days, we find double positive (9% mCherry^+^ and BFP^+^) cells as desired (**Fig. 5E**), without any reduction in mCherry, showing a second gene can be delivered without losing the first cargo. To confirm Cp36 is unidirectional, we used a plasmid recombination assay and saw no recombination between attL and attR (**Fig. S5F, G**).

Finally, we demonstrate two other multi-targeting orthologs with efficiencies of 13% and 35% (**Fig. S5H**), demonstrating this multi-targeting clade is a rich repository of efficient recombinases. These results reveal the existence of a subset of LSRs, not previously tested in eukaryotic cells, with highly efficient unidirectional integration activity and higher specificity compared to transposase systems.

## DISCUSSION

DNA-targeting enzymes derived from the microbial diversity have revolutionized molecular biology and genome engineering. Due to their ability to transfer large DNA cargo, integrase systems such as recombinases and transposases have been commonly employed for workflows such as Gateway cloning or generation of stable cell lines (Sandoval-Villegas et al. 2021; X. Wang et al. 2017). Despite longstanding efforts to adapt them for genome editing, the low efficiency and small number of known large serine recombinases have greatly limited their broader utility for mammalian engineering (Merrick, Zhao, and Rosser 2018). We sought to address these challenges by automatically processing a large number of microbial mobile genetic elements in order to identify novel LSR enzymes. By increasing the number of known LSR and cognate attachment site combinations by >100-fold relative to previous work (Yang et al. 2014), we identified key functional classes of LSRs based on their ability to target the human genome. In summary, we identify and experimentally validate LSRs in three major classes with potential clinical and research utility (**Table S1**).

First, we identified an array of new landing pad LSRs, which outperform the wild-type Bxb1 by 2 to 7-fold in episomal and chromosomal integration efficiency. In addition to facilitating single payload insertions at efficiencies of 40-70% without selection, these landing pad LSRs can be programmed to direct cargo DNA into specific landing pad sites based on matching of the core dinucleotide in the attachment sites, suggesting combinations of donors could be specifically addressed to an array of landing pads in the same cell (Ghosh, Kim, and Hatfull 2003). We further demonstrate the utility of landing pad LSRs for sensitive functional genomics applications, such as parallel reporter assays in the endogenous genome. These results expand our capability to build efficient and specific landing pads in human cells and other organisms for applications in research.

Another highly desirable LSR application would be to identify a variant that can achieve both efficient and site-specific integration in the human genome without requiring a landing pad. Despite extensive characterization of PhiC31 with the goal of therapeutic gene integration (Calos 2006), its integration rate does not surpass 3% across at least 42 pseudosites (Thyagarajan et al. 2008; Jaichandran Sivalingam et al. 2010; Chalberg et al. 2006). We reasoned that we could leverage our LSR database by computationally searching for matches between native attachment sites and the human genome sequence. Excitingly, we demonstrate that we could predict integration sites using BLAST homology of their attachment sites with the human genome. Candidates tested so far target more than a single site in the human genome, which is many-fold larger than the average bacterial genome, but we identified LSRs that integrate into their top site at frequencies from 6% (Sp56, Enc3) to 34.3% (Pf80) of all integrations. This is much more specific than other microbial integrases in human cells like PiggyBac transposase (Wilson, Coates, and George 2007). Taken together, we demonstrate the ability to identify site-specific LSRs by computationally comparing native attachment sites with a target genome. Our LSR database also includes candidates that could directly target non-human genomes, including plants, microbes, and model- or non-model organisms in which stable transgenesis is currently difficult.

Because our computational approach identifies candidate LSRs as well as their target sites, this expansive database provides insight into the innate targeting specificity of a given LSR. Some appear to target unique sites in bacteria while others are more promiscuous, a group that we describe as multi-targeting. When we introduced our multi-targeting LSR Cp36 into human cells, we found that it integrated cargo DNA into the human genome with high efficiency (>40%) at multiple sites. Cp36 compares favorably with the Super PiggyBac transposase in efficiency, unidirectionality (it does not excise its previous insertions when re-used), and design (it only requires appending a short attP to one end of its cargo rather than 200-300 nt flanking arms). Super PiggyBac has been extensively used for transgenesis, mutagenesis, and for therapeutic purposes such as CAR T cell engineering (Yusa 2015; X. Li et al. 2013; Ptáčková et al. 2018). We envision multi-targeting LSRs supplanting transposases and retroviruses in many applications that require high efficiency integration with better defined target sites, such as cell therapies.

Interestingly, while LSRs alone perform a unidirectional integration reaction, they can perform the reverse excision reaction when co-expressed with a reverse directionality factor (RDF) protein (Khaleel et al. 2011; Fogg et al. 2018). An exciting future direction is to extend the bioinformatic search of these mobile genetic elements in order to retrieve the RDF corresponding to each of these LSRs. Such RDFs could expand LSR utility for synthetic biology and, potentially, provide a form of antidote or safety switch in cases where an LSR-mediated integration needs to be removed.

A potential limitation of LSRs is that they are not readily reprogrammed to target new sequences. Promisingly, we find natural attachment sites vary widely across LSR clusters, suggesting arbitrary sequences could be targetable by LSRs. Further work to dissect LSR structure-function relationships with their target DNA sequence could enable the design of synthetic LSRs that can be reprogrammed to target new locations in genomes, providing a simple single effector protein tool to integrate large cargoes into arbitrary locations. In addition, LSRs or domains of LSRs could potentially be combined with programmable CRISPR targeting system to generate an approach that combines both site specificity and host-independent DNA recombination. While this manuscript was in preparation, such an approach was described with prime editors (Ioannidi et al. 2021; Anzalone et al. 2021). An exciting future direction would be to combine prime editors with the efficient and specific landing pad LSRs described here to most efficiently integrate large cargos into programmed locations.

Taken together, we envision diverse applications of integrase systems for reliable, stable, and unidirectional targeting of the genome, such as functional genomics screens where controlled insertion of unique library elements into unique single cells is desired (Matreyek et al. 2020). Second, these landing pads could be useful in the development of engineered cells or cellular therapies, where custom combinations of genes can be introduced to induce cell type-specific differentiation or to control cell behavior via synthetic gene circuits. Third, both endogenous and synthetically inserted LSR attachment sites can be used as scratchpads to record and reconstruct cell lineages during cell fate specification in development or disease models (Chow et al. 2021). Finally, these LSRs could also enable larger scale genome engineering, including controlled models of large structural rearrangements by installing the attachment sites at distal sites in a genome (Esvelt and Wang 2013). Beyond LSRs, there are many more DNA mobilization genes lying in wait within massive sequence databases, providing an expansive opportunity to derive insights into their mechanisms of protein-DNA interaction and enrich the genome engineering toolbox.

## FIGURES

**Figure S1.**
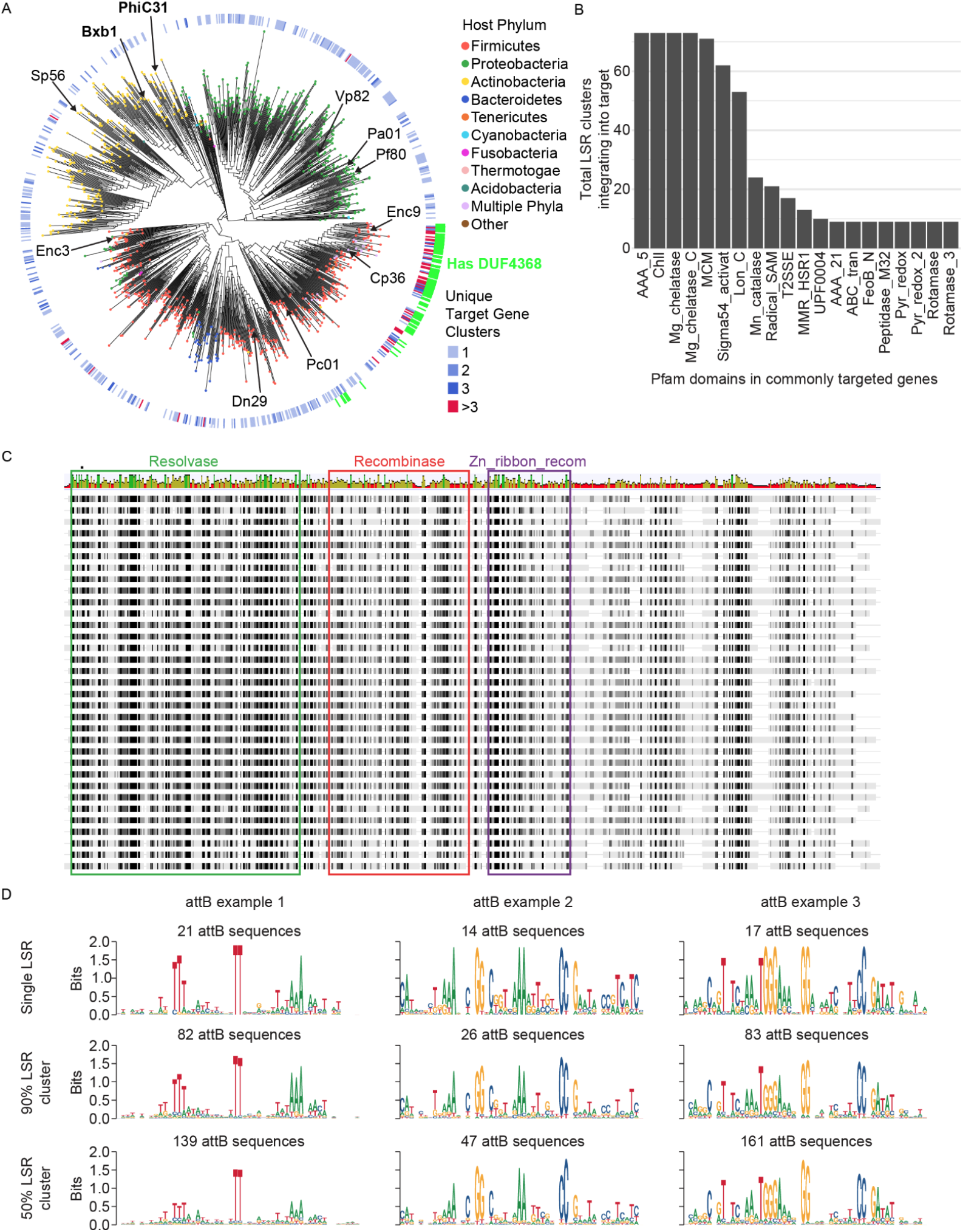
Bioinformatic discovery of large serine recombinases and their target attachment sites. **A**. Phylogenetic tree of 1,081 LSR clusters (50% identity). Tips are colored according to the phylum of bacterial host species. First heat map ring is colored according to the number of unique target gene clusters that each LSR cluster is predicted to integrate into, as in **Fig. 1B**. The second ring of green annotations indicate LSR clusters that are predicted to contain the DUF4368 Pfam domain. Clusters containing Bxb1 and PhiC31 are indicated in bold text, and clusters for select candidates with experimental validation are also indicated. **B**. Histogram of Pfam domains most commonly found in target genes. Each target gene was annotated using Pfam HMM models, and then the total number of LSR clusters that integrate into genes containing each Pfam domain was calculated. **C**. Alignment of LSR sequences that are presented in **Fig. 1F**. Pfam domains are indicated on top. Above each aligned amino acid position, the height and color of each bar indicates the mean pairwise identity over all pairs in the column, with green indicating 100% identity across all sequences, green-brown indicating above 30% identity and below 100% identity, and red indicating below 30% identity. **D**. Examples of predicted attB motifs. Each column represents a different LSR attB motif. The first row shows motifs that were derived from different attB sequences that were all targeted by a single, unique LSR protein. The second row shows motifs that were derived from attB sequences that were targeted by LSR proteins that fell into a single 90% identity cluster. The third row shows motifs that were derived from attB sequences that were targeted by LSR proteins that fell into a single 50% identity cluster.

**Figure S2.**
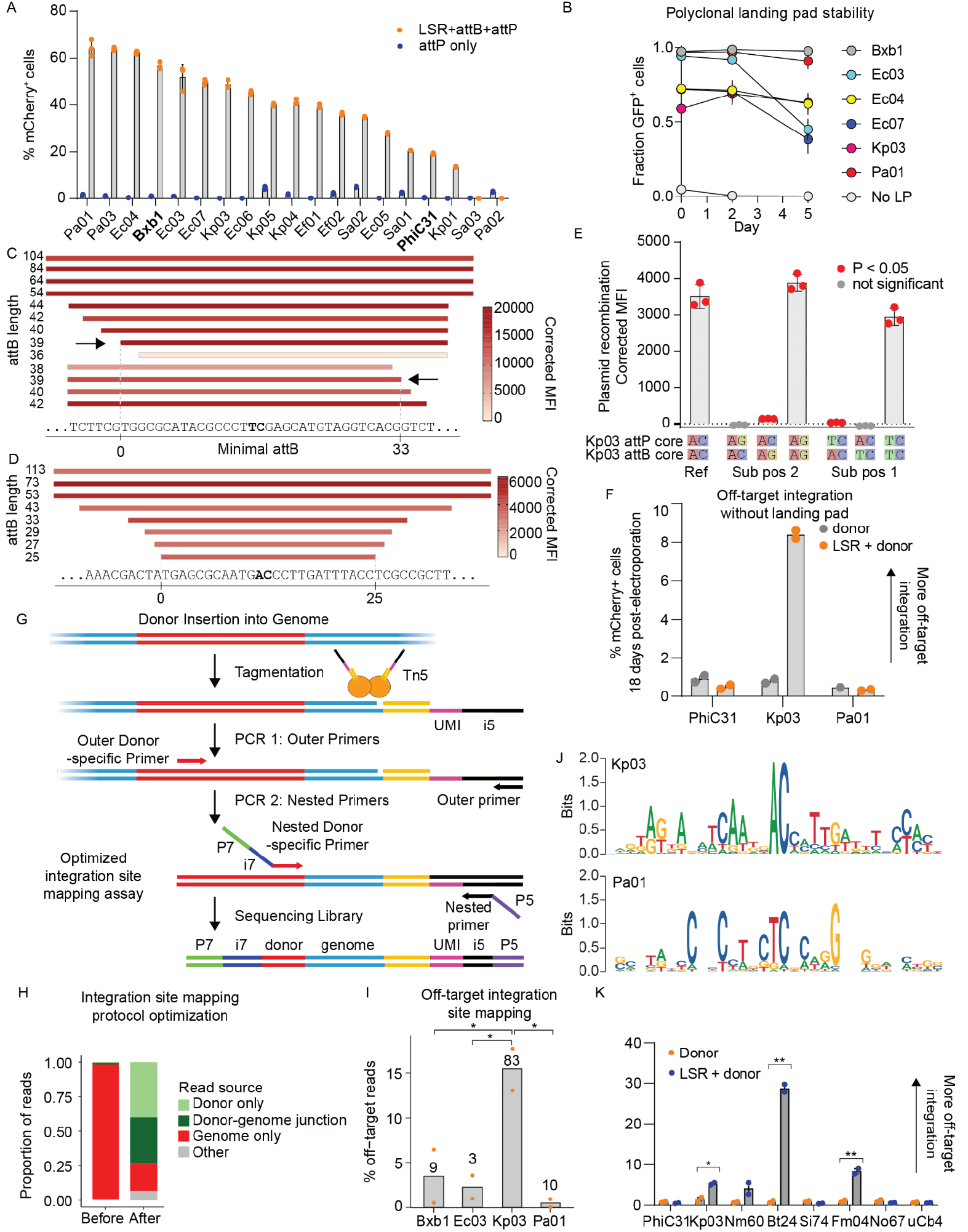
New landing pad LSRs have short attachment sites, can be multiplexed by core swaps, and can be highly specific. **A**. Plasmid recombination assay of predicted LSRs and att sites in HEK293FT cells, shown as percentage of mCherry^+^ cells gated on GFP positive cells. mCherry and GFP gating is determined based on an empty backbone transfection. Dots show each transfection replicate, error = SD (n=3 transfection replicates). **B**. Stability of polyclonal landing pads expressing LSR-GFP as measured by flow cytometry over time. These cells are not electroporated with donor and day 5 was the same day of measurement as for **Fig. 2F** (n=2 independently transduced biological replicates). **C**. Minimization of Pa01 attB sequence by trimming nucleotides from either end and using the plasmid recombination assay. Arrows indicate shortest attB which did not disrupt recombination activity. The inferred 33 bp minimal attB as determined by this experiment is shown between vertical lines at the bottom. Colored rectangles show mean corrected mCherry MFI (n=3 transfection replicates in HEK293FT cells). The attB in the top rectangle extends in both directions and is the full length attB as retrieved from the LSR database and used in **Fig. 2B and 2C**. A predicted dinucleotide core as determined by off-target integration site mapping is highlighted in bold. **D**. Minimization of Kp03 attB sequence by trimming nucleotides from both ends using the plasmid recombination assay. The shortest tested attB, which efficiently recombined, was 25 nucleotides suggesting the minimal attB is 25 nucleotides or shorter. Colored rectangles show mean mCherry MFI normalized to attD only MFI (n=3). The attB in the top rectangle extends in both directions and is the full length attB as retrieved from the LSR database and used in **Fig. 2B and 2C**. The dinucleotide core, as determined by core swapping experiments in (G) and off-target integration site mapping, is shown in bold text. **E**. Kp03 dinucleotide core substitution in the plasmid recombination assay. AC is the native dinucleotide core sequence. Values are mean ± SD with n=3 transfection replicates in HEK293FT cells. P-value determined by one-tailed t-test. **F**. Flow cytometry measuring mCherry^+^ cells 18 days after LSR and donor co-electroporation into WT K562 cells that lack a landing pad. attD donor contains its own EF-1α promoter and attD donor-only is a negative control (n=2 transfection replicates). **G**. Schematic of optimized integration site mapping assay, a modified version of UDiTaS (Giannoukos et al. 2018). Addition of a round of amplification using a nested donor primer is expected to enrich for desired target-derived reads, which includes both donor-only reads and donor-genome junction reads (see Methods for details). **H**. Proportion of reads derived from different sources in the integration site mapping assay. On the left, the proportions before assay optimization, and after optimization on the right. Both runs are of Cp36 circular donor experiments, but in two different cell types (HEK293FT on the left, K562 on the right). Target-derived reads are those that come from the donor only (light green) or the donor-genome integration junction reads (dark green). **I**. Genome-wide integration site mapping by next generation sequencing to measure the percentage of reads found in the genome outside the expected landing pad. Showing raw (non-unique) reads found at off-targets as a percentage of all reads (* = P < 0.05, one-tailed t-test). For Kp03, Ec03, and Pa01, n = 2 independent clonal landing pad lines with maximal mCherry 11 days post donor electroporation. For Bxb1, showing two technical replicates of a single clonal landing pad line with maximal mCherry 11 days post donor electroporation. Numbers near the top of each bar indicate the total number of off-target loci. **J**. Showing a target site motif of the top 25 most frequent human genome off-target sites for landing pad candidates Kp03 (top) and Pa01 (bottom). Core dinucleotides are strongly conserved among integration sites for both candidates. **K**. Flow cytometry measuring mCherry^+^ cells 18 days after LSR and donor co-electroporation into WT K562 cells that lack a landing pad. The donor plasmid contains its own EF-1α promoter driving mCherry expression (* = P < 0.05, ** = P < 0.005, one-tailed t-test). (n = 2 transfection replicates, error = SEM)

**Figure S3:**
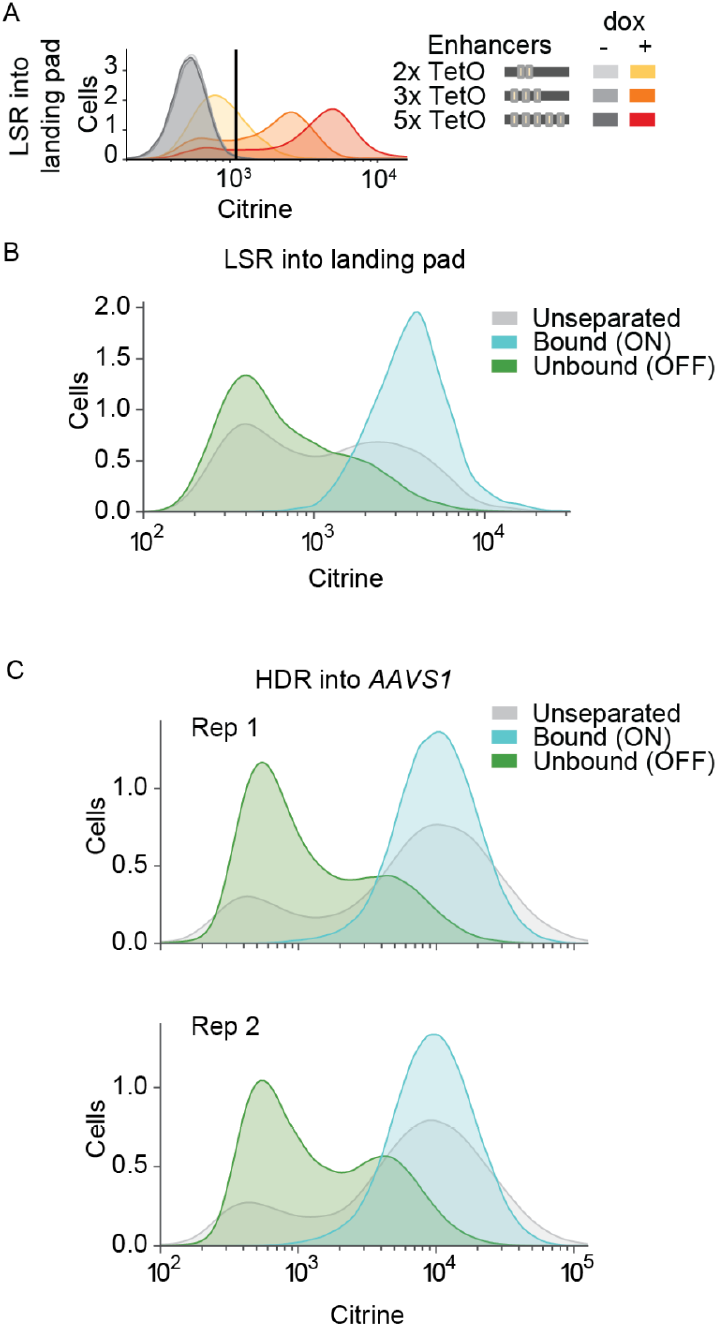
Parallel reporter assay using magnetic separation. **A**. Flow cytometry measurements of the landing pad citrine reporter 2 days after induction with doxycycline, in a distinct replicate from the experiment shown in **Fig. 3B**. This replicate is time-matched with the LSR landing pad PRA shown in **Fig. 3C**, wherein reporters were delivered, selected with puromycin for 8 days, grown for 2 weeks, and then induced with doxycycline for 2 days before analysis. Vertical line marks the linear gate for Citrine^+^ cells, which are shown in **Fig. 3C** (n=1 cell line replicate). **B**. Efficiency of magnetic separation for LSR landing pad PRA cells, corresponding to the samples sequenced and shown in **Fig. 3C**. Unseparated sample is the pooled, dox-induced cells before mixing with magnetic beads. **C**. Efficiency of magnetic separation for HDR-integrated PRA cells, corresponding to the samples sequenced and shown in **Fig. 3C**.

**Figure S4.**
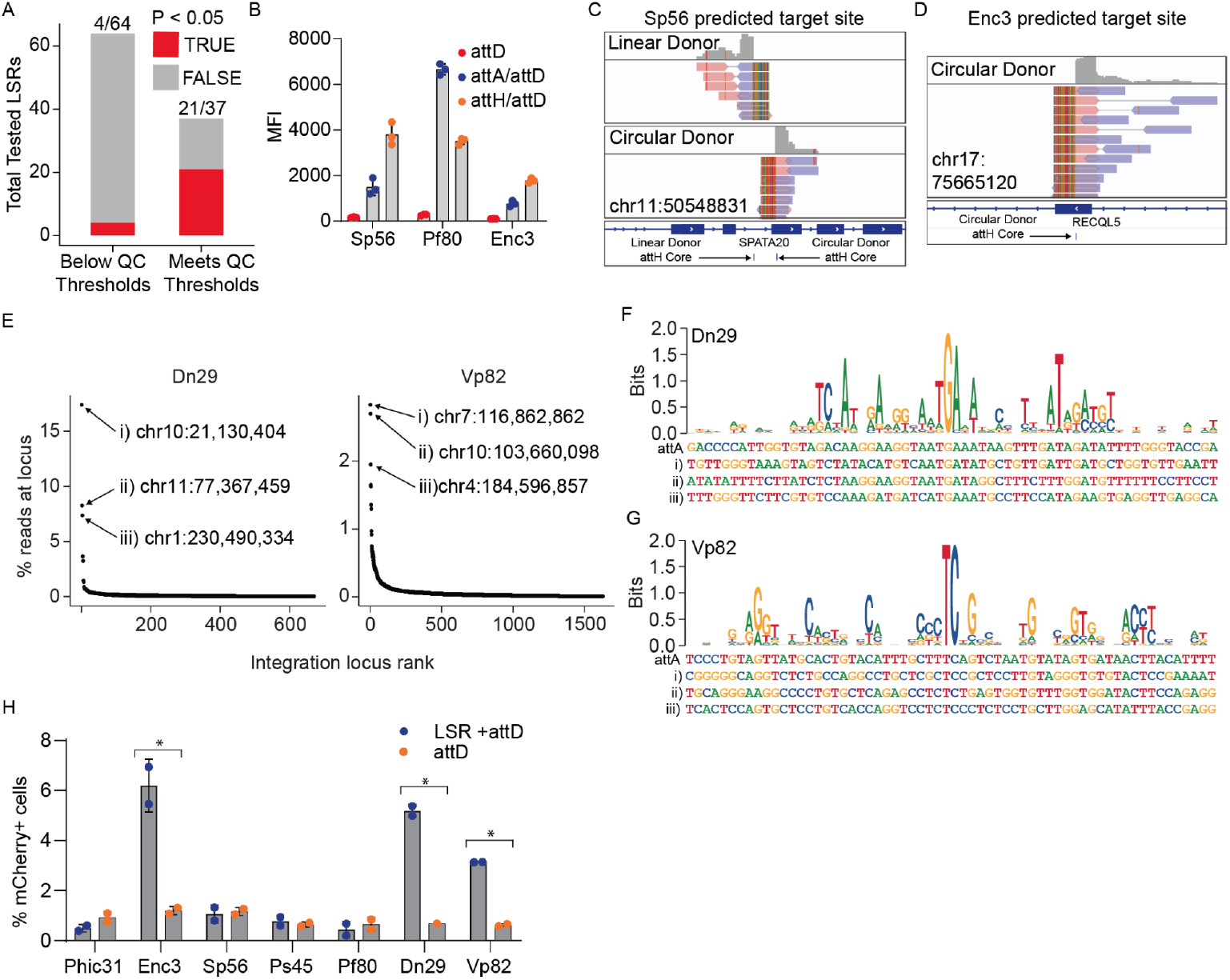
Human genome-targeting recombinases target specific and predictable sites. **A**. Proportion of genome-targeting LSR candidates that mediate significant recombination in the plasmid recombination assay with and without application of quality control (QC) thresholds for LSR candidate selection. The numbers above each bar indicate the (number of candidates that met P < 0.05 in the plasmid recombination assay) / (total number of tested candidates). **B**. Plasmid recombination assay for top genome-targeting candidates using predicted attH sites. Showing that recombination between plasmids does occur between predicted attachment sites and their human genome target sequences. (n=3 transfection replicates) **C**. Same as in **Fig. 3E**, but for Sp56. The orientation and location of the integration changes when using a linear donor, whereas the exact predicted integration site is targeted with a circular donor. **D**. Same as (C), but for the predicted target site of Enc3. **E**. Integration site mapping results for Dn29, and Vp82. Top 3 targeted human genome sites are labeled in each panel. The most commonly targeted site for Dn29 accounts for 17.4% of unique reads mapping to integration sites, suggesting that this candidate has a favorable mix of efficiency and specificity. **F**. Sequence logo depicting the target site motifs for Dn29, built using the top 25 human genome target sites, ranked according to the number of reads at each site. Giving examples of complete genomic target sites beneath the motif, as specified in (E). **G**. Same as (F), but showing the target site motifs for Vp82. **H**. Human genome integration efficiency assay results of the top candidates. PhiC31 is a previously known genome targeting LSR used as a control, although its efficiency is below the limit of detection (∼1% of cells). Bars are mean, dots are individual transfections. Error = s.d. (*=P<.05, one-tailed t-test)

**Figure S5.**
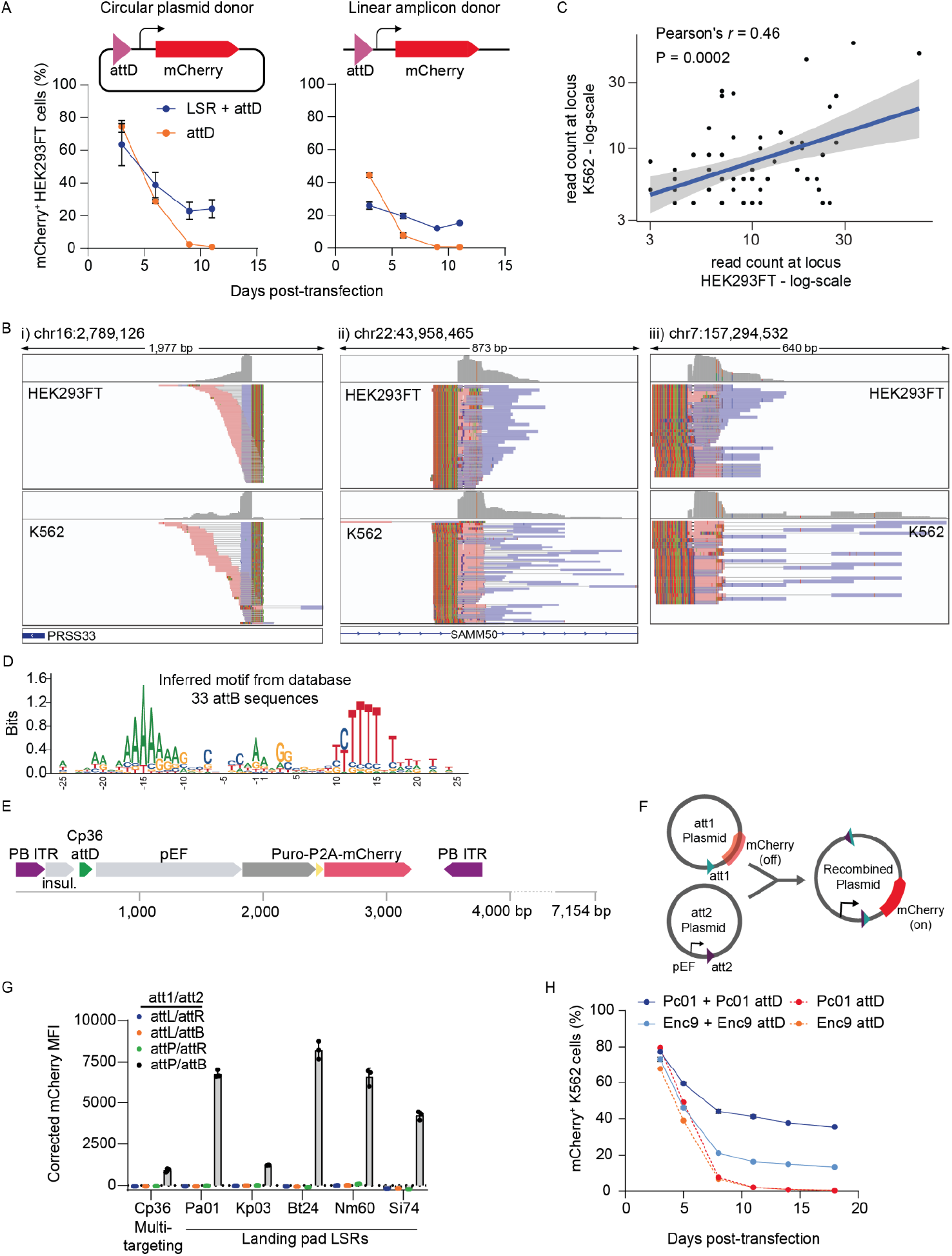
Multi-targeting recombinases are efficient and unidirectional integrases. **A**. Cp36 co-transfected with either a circular plasmid or linear amplicon attD-pEF-1α-mCherry donor in HEK293FT cells, compared with attD-only control (n = 2 transfection replicates), error = SEM. Stable fluorescence in cells treated with Cp36 was measured by flow cytometry. **B**. Integration site mapping shows precise integration into the same sites in multiple genomic DNA fragments from cells post-transfection with Cp36, suggesting these are recurring hotspots. Aligned reads colored according to forward strand (red) or reverse strand (blue), with paired reads joined by a black line. Soft-clipped portions of the aligned reads colored to show where the read crosses over from the human genome into the Cp36 donor sequence, and the reference genome sequence on the bottom. Showing the top three integration sites specified in **Fig. 5C**. At the integration sites found at locus i and ii, we see reads in both the forward and reverse orientation, suggesting both integration orientations are possible at these sites, although one orientation dominates. **C**. Correlation between read counts from the Cp36 integration site mapping assay across HEK293FT and K562 cell lines. Showing the top 61 shared loci, all of which are found among the top 200 most frequently targeted sites in the two cell types. **D**. Showing target site motif as predicted using 33 attB sequences in the LSR-attachment site database that are targeted by LSRs that fall in the same 50% amino acid identity cluster as Cp36. Method used to construct this motif is the same as in **Fig. 1H** and **Fig. S1D. E**. Schematic of donor plasmid used for direct comparison of Cp36 and PiggyBac that contains both the PB inverted terminal repeats (ITRs) and the Cp36 attD. **F**. Schematic of plasmid recombination assay to determine directionality of LSR recombination. In addition to the original attP/attB, attL/attR were also used in various combinations with attP/attB. If significant mCherry^+^ fluorescence is detected in the attL/attR experiment, it would imply that the LSR was bi-directional, being capable of excision. The LSR is co-transfected on a third plasmid. **G**. Results of plasmid recombination assay to determine LSR directionality. Showing mCherry MFI, corrected according to a control that lacked the LSR plasmid. Cp36 and various landing pad LSR candidates were tested. Results indicate that recombination is unidirectional, as significant mCherry fluorescence is only detected in the attP/attB condition. Bars are mean, dots are transfection replicates. Error=s.d. **H**. Additional multi-targeting LSR candidates validated using the pseudosite integration assay. Showing two additional candidates, Pc01 and Enc9, which are considered multi-targeting, with Pc01 containing DUF4368 and residing in a clade that is closely related to the primary multi-targeting clade, and Enc9 residing directly in the primary multi-targeting clade shown in **Fig. 1B** and **Fig. S1A**. Dots are mean, Error = s.d. (n=2 electroporation replicates).

## METHODS

### Cell lines and cell culture

Experiments were carried out in K562 cells (ATCC CCL-243) and HEK-293FT cells. K562 cells were cultured in a controlled humidified incubator at 37°C and 5% CO_2_, in RPMI 1640 (Gibco) media supplemented with 10% FBS (Hyclone), penicillin (10,000 I.U./mL), streptomycin (10,000 ug/mL), and L-glutamine (2 mM). HEK-293FT cells, as well as HEK-293T and HEK-293T-LentiX cells used to produce lentivirus, as described below, were grown in DMEM (Gibco) media supplemented with 10% FBS (Hyclone), penicillin (10,000 I.U./mL), and streptomycin (10,000 ug/mL).

### Computational workflow to identify thousands of LSRs and cognate attachment sites

The LSR-identification workflow was implemented as described schematically in **Fig. 1A**. 146,028 bacterial isolate genomes available in the NCBI RefSeq database were identified on August 22nd, 2019. Genomes were then clustered at the species level using the NCBI taxon ID and the TaxonKit tool (Shen & Xiong, 2019). Genomes within each species were randomized and batched into sets of 50 and 20 genomes, where the first batch included 50 genomes and all subsequent batches contained 20 genomes. Each batch was then processed by downloading all relevant genomes from NCBI, annotating coding sequences in each genome with Prodigal (Hyatt et al., 2010), and then searching for all encoded proteins that contained a predicted Recombinase Pfam domain using HMMER (El-Gebali et al., 2019; HMMER, n.d.). Genomes that contained a predicted LSR were then compared to genomes that lacked that same LSR using the *MGEfinder* command *wholegenome*, which was developed for this purpose by adapting the default *MGEfinder* to work with draft genomes. If MGE boundaries that contained the LSR were identified, all of the relevant sequence data was saved and stored in a database. The workflow was parallelized using Google Cloud virtual machines.

After this initial round of LSR discovery was complete, a modified approach was taken to further expand the database and avoid redundant searches. First, bacterial species with a high number of isolate genomes available in the first round were analyzed to determine if further inspection of these genomes would be necessary. Rarefaction curves representing the number of new LSR families identified with each additional genome analyzed were estimated for these common species, and species that appeared saturated (i.e. less than 1 new cluster per 1000 genomes analyzed) were considered “complete,” meaning no further genomes belonging to this species would be analyzed. Next, 48,557 genomes that met these filtering criteria were downloaded from the GenBank database and prepared for further analysis. The analysis was very similar to round 1, but with some notable differences. First, a database of over 496,133 isolate genomes from the RefSeq and GenBank genomes was constructed. PhyloPhlAn marker genes were then extracted from all of these genomes (Asnicar et al. 2020). Next, for each genome that was found to contain a given LSR, closely related isolates found in the database were selected according to marker gene homology and used for the comparative genomics analysis and further LSR discovery. This marker gene search approach was made available in a public Github repository (https://github.com/bhattlab/GenomeSearch). This second round of LSR and attachment site discovery increased the total number of candidates by approximately 32%.

### Predicting LSR target site specificity

LSR sequences were clustered at 90% and 50% identity using MMseqs2 (Hauser, Steinegger, and Söding 2016). Protein sequences that overlapped with predicted attachment sites were extracted from their genome of origin and clustered with all other target proteins at 50% identity using MMseqs2. LSR-attachment site combinations that were found to meet quality control filters were considered. To identify site-specific LSRs, only LSRs clustered at 50% identity and target genes clustered at 50% amino acid identity were considered. Next, LSR-target pairs were filtered to only include target gene clusters that were targeted by 3 or more LSR clusters. Next, only LSR clusters that targeted a single target gene cluster were considered. The remaining sets of LSR clusters were considered to be single-targeting, meaning that they were believed to site-specifically target only one gene cluster. Multi-targeting, or transposable LSRs with minimal site-specificity, were identified. Only LSRs clustered at 90% identity and target genes clustered at 50% amino acid identity were considered. Next, the total number of target gene clusters that were targeted by each LSR cluster were counted, and LSR clusters that targeted only one gene cluster were removed from consideration. Next, the remaining LSRs were binned according to the number of protein clusters that they targeted. For the purposes of this paper, “>3” are considered fully multi-targeting. Each 50% identity LSR cluster was then assigned to a multi-targeting bin according to the highest bin attained by any one 90% LSR cluster found within the 50% identity LSR cluster.

### Phylogenetic tree construction

Representative sequences of each quality-controlled 50% identity LSR cluster were used to construct the phylogenetic tree. LSRs were aligned using MAFFT in G-INS-i mode (Katoh and Standley 2013), and IQ-TREE was then used to generate a consensus tree using 1000 bootstrap replicates and automatic model selection.

### Phylogenetic analysis of site-specific integrases targeting a conserved attachment site

One example of several site-specific integrases targeting a conserved attachment site is shown in **Fig. 1E**. All attB attachment sites were clustered at 80% identity using MMseqs2 (Steinegger and Söding 2017). Candidates were filtered to include only those that met QC thresholds, and then attB sites that were ranked by the number of LSR clusters that were found to target them. An example attB cluster was chosen for further analysis. All LSRs that targeted this attB cluster were extracted from the database, and were aligned using the MAFFT-LINSI algorithm (Katoh and Standley 2013). Amino acid identity distances between all LSRs were calculated, and the distance matrix was used to create a hierarchical tree in R. LSRs that were 99% identical at the amino acid level or more were collapsed into a single cluster. This hierarchical tree was visualized and shown in **Fig. 1E**, along with all attB sites that were targeted by the LSRs.

### Identifying target site motifs from attachment sites in the LSR database

The attachment sites associated with multi-targeting LSRs in the database could be analyzed to determine target site motifs, as shown in **Fig. 1H** and **Fig. S1D**. Multi-targeting LSRs in the database were analyzed at the level of individual proteins, at the level of 90% amino acid identity clusters, and at the level of 50% amino acid identity clusters. For each of these levels, only candidates that were found to target more than 10 unique attB sequences or 10 target genes clustered at 50% amino acid identity were kept. Then all of the corresponding attB sequences were extracted, with only one attachment site per target gene cluster being extracted to avoid redundancy. These attB sequences were then initially aligned using MAFFT-LINSI (Katoh and Standley 2013). Next, possible core dinucleotides were identified in each alignment by extracting all dinucleotides in the alignment, and ranking them by the conservation of their most frequent nucleotides and their proximity to the center of the attB sequences, using a custom score that equally weighted high nucleotide conservation and normalized distance to the attB center. Candidates were then re-aligned only with respect to these predicted dinucleotide cores, rather than using an alignment algorithm such as MAFFT. These alignments were then visualized in using ggseqlogo to identify conserved target site motifs (Wagih 2017).

### Initial landing pad LSR candidate selection

LSRs for the initial set of 17 landing-pad candidates were identified by searching for the Recombinase Pfam domain among the MGEs we previously identified (Durrant et al., 2019; El-Gebali et al., 2019). The identity of the attachment site was inferred from the boundaries of the MGE that contained each LSR. For example, imagine a sequence nucleotides that have the following structure:

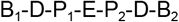

Where B_1_ indicates the sequence flanking the MGE insertion on the 5’ end, D indicates the target site duplication created upon insertion (if it exists), P_1_ indicates the sequence flanking the 5’ integration boundary that is included in the MGE, E is the intervening MGE, P_2_ indicates the sequence flanking the 3’ integration boundary that is included in the MGE, and B_2_ indicates the sequence flanking the MGE insertion on the 3’ end, then the attB and attP sequences can be reconstructed as:

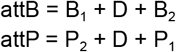

Where the “+” operator in this case indicates nucleotide sequence concatenation.

Candidates were then annotated to determine features such as: 1) Whether or not the element was predicted to be a phage element (Arndt et al., 2016), 2) how many isolates contain the integrated MGE, and 3) how often MGEs containing distinct LSRs will integrate at the same location in the genome. Candidates were then given higher priority if they were contained within predicted phage elements, if they appeared in multiple isolates, and if the attachment sites were targeted by multiple distinct LSRs. A final list of 17 candidates, listed in **Fig. 2B**, was then taken forward and validated experimentally.

### Subsequent selection of LSR candidates of high quality

As subsequent batches of LSRs were ordered and tested in our various assays, we improved our quality control criteria for selecting further candidates to synthesize and assay. In our initial batch of human genome-targeting candidates, few quality control filters were put in place. Subsequent batches were more stringently quality controlled. We settled on one set of quality control criteria that dramatically increased experimental validation rate. First, LSRs with large attachment site centers, above 20 base pairs in length, were removed. The attachment site center is the portion of the attB and the attP that are identical, and should contain the dinucleotide center. Next, LSRs with attachment sites with more than 5% of their nucleotides being ambiguous in the original genome assemblies were removed. Next, only LSRs between 400 amino acids and 650 amino acids were kept. Next, only predicted LSRs that contained at least one of the three main LSR Pfam domains were retained (Resolvase, Recombinase, and Zn_ribbon_recom). Next, LSRs were removed from consideration if their sequences contained more than 5% ambiguous amino acids. Next, only LSRs that were found on integrative mobile genetic elements that were less than 200 kilobases in length were retained, where larger elements were presumed to be technical artifacts. And finally, only LSRs that were within 500 nucleotides of their predicted attachment sites were retained. Candidates that met all of these filters were considered to meet quality-control thresholds.

### Plasmid recombination assay to validate LSR-attD-attA predictions

Three plasmids were designed for each LSR candidate to test recombination function on an episomal reporter. The effector plasmid contains the EF-1α promoter, followed by the recombinase coding sequence (codon optimized for human cells), a 2A self-cleaving peptide, and an EGFP coding sequence. The attA plasmid contains an EF-1α promoter, followed by the attA sequence, followed by mTagBFP2 coding sequence, which should constitutively express the mTagBFP2 protein in human cells. The attD plasmid includes only the attD sequence followed by the mCherry coding sequence, which should produce no fluorescent mCherry prior to integration. 20,000 HEK-293FT cells were plated into 96 well plates and transfected one day later with 200 ng of effector plasmid, 70 ng of attA plasmid, and 50 ng of attD plasmid using Lipofectamine 2000 (Invitrogen). 2-3 days after transfection of cells with all three plasmids, cells were then measured using flow cytometry on an Attune NxT Flow Cytometer (ThermoFisher). HEK-293FT cells were lifted from the plate using TrypLE (Gibco), and resuspended in Stain Buffer (BD) before flow. These experiments were conducted in triplicate transfections. Cells were gated for single cells using forward and side scatter, and then on cells expressing fluorescent EGFP. Next, mTagBFP2 fluorescence was measured to indicate the amount of un-recombined attD plasmids, and mCherry fluorescence was measured to indicate the amount of recombinant plasmid. Corrected mean fluorescent intensity (MFI) is the MFI after subtracting the average MFI of all matching attD-only control replicates. mCherry and EGFP gating was determined based on an empty backbone transfection.

An experiment testing recombinases with matched and unmatched attB and attP plasmids was performed similarly, in K562 cells. 1.2×10^6^ K562 cells were electroporated in 100 μl Amaxa solution (Lonza Nucleofector 2b, program T-016), with 300 ng of the 11.6 kb LSR plasmid, 869 ng of the 4.2 kb attB plasmid, and 621 ng of the 3 kb attP plasmid. 3 days after transfection, mCherry MFI of ungated cells was measured by flow cytometry on a BD Accuri C6 cytometer.

### Landing pad cell line production

Landing pad LSR candidates were cloned into lentiviral plasmids under the expression of the strong pEF-1α promoter, with their attB site in between the promoter and start codon, and with a 2A-EGFP fluorescent marker downstream the LSR coding sequence. Lentivirus production and spinfection of K562 cells were performed as follows. We plated HEK-293T cells on 6-well tissue culture plates. In each well, 5×10^5^ HEK-293T cells were plated in 2 mL of DMEM, grown overnight, and then transfected with 0.75 μg of an equimolar mixture of the three third-generation packaging plasmids (pMD2.G, psPAX2, pMDLg/pRRE) and 0.75 μg of LSR vectors using 10 μl of polyethylenimine (PEI, Polysciences #23966) and 200 μl of cold serum free DMEM. pMD2.G (Addgene plasmid #12259; http://n2t.net/addgene:12259; RRID:Addgene_12259), psPAX2 (Addgene plasmid #12260; http://n2t.net/addgene:12260; RRID:Addgene_12260), and pMDLg/pRRE (Addgene plasmid #12251; http://n2t.net/addgene:12251; RRID:Addgene_12251) were gifts from Didier Trono. After 24 hours, 3 mL of DMEM was added to the cells, and after 72 hours of incubation, lentivirus was harvested. We filtered the pooled lentivirus through a 0.45-μm PVDF filter (Millipore) to remove any cellular debris.

To create polyclonal landing pad cell lines, 2 mL of lentiviral supernatant and 8 μg/ml polybrene was used on 3×10^5^ K562 cells to ensure a high multiplicity of infection. These cells were infected by spinfection for 30 minutes at 1000 x g at 33°C followed by overnight infection. The next day, the cells were spun down and resuspended in fresh media. This resulted in >50% EGFP^+^ cell populations, suggesting each cell likely contained multiple landing pad copies. To create clonal landing pad cell lines, lentivirus doses of 50, 100, and 200 μl were used for each vector, in order to find a condition with low multiplicity of infection wherein each transduced cell would be likely to contain only a single integrated copy of the landing pad. 3×10^5^ K562 cells were mixed with the lentiviruses in 8 μg/ml polybrene and infected overnight, without spinfection. Infected cells grew for 3 days and then infection efficiency was measured using flow cytometry to measure EGFP (BD Accuri C6); the dose that gave rise to 5 - 15% EGFP^+^ cells was selected for each LSR for further experiments. Ten days later, these EGFP^+^ cells were sorted into a 96-well plate with a single cell in each well, in order to derive clonal lines with a single landing pad location. Two weeks later, 4 clones for each LSR with a unimodal high EGFP expression level were selected for expansion and subsequent experiments.

### Landing pad integration efficiency assay

Landing pad cell lines were electroporated in 100 μL Amaxa solution (Lonza Nucleofector 2b, program T-016) with the promoterless mCherry donor containing the matching attP at a dose of either 1000, 2000, or 5000 ng donor plasmid. At timepoints from 5 - 12 days post-electroporation, the cells were subjected to flow cytometry to measure mCherry and EGFP (BD Accuri C6). In one experiment, measurements were made on a BioRad ZE5 cytometer, which is noted in the associated figure legend.

### Pseudosite integration efficiency assay to measure integration percent into the WT genome

To determine the percentage of integration of attD donors into pseudosites in the human genome, attD sequences were cloned into a plasmid containing an pEF-1α promoter followed by mCherry, a P2A self-cleaving peptide, and a puromycin resistance marker. Integration efficiency was measured in both K562 and HEK-293FT cells. In K562 cells, 1.0×10^6^ cells were electroporated in 100 μL Amaxa solution (Lonza Nucleofector SF, program FF-120), with 3000 ng LSR plasmid and 2000 ng pseudosite attD plasmid. As a non-matching LSR control, 3000 ng of Bxb1 was substituted for the correct LSR plasmid. The cells were cultured between 2×10^5^ cells/mL and 1×10^6^ cells/mL for 2-3 weeks.

In HEK-293FT cells, 20,000 cells were plated into 96 well plates, and transfected a day later with 200 ng of LSR plasmid and 178 ng of pseudosite attD plasmid using Lipofectamine 2000 (Invitrogen). As a non-matching LSR control, 200 ng of Bxb1 was substituted for the correct LSR plasmid. Additionally, a linear version of the pseudosite attD donor was also tested for integration activity in HEK-293FT cells. To create the linear donors, pseudosite attD plasmids were PCR amplified using the KAPA Hifi HotStart ReadyMix (Roche), amplifying the pEF-1α promoter followed by mCherry, a P2A self-cleaving peptide, and a puromycin resistance marker and removing the plasmid bacterial elements. The PCR product was gel extracted with the Monarch DNA Gel Extraction Kit (NEB). 20,000 HEK-293FT cells were plated into 96 well plates, and transfected a day later with 300 ng LSR plasmid and 24 ng of the linear pesudosite attD donor. As a non-matching LSR control, 300 ng of Bxb1 was substituted for the correct LSR plasmid.

For all K562 and HEK-293FT transfections, 100 uL of each sample was run on the Attune NxT Flow Cytometer every 3-4 days to measure the mCherry signal. After 2-3 weeks, transiently transfected plasmid was nearly fully diluted out in the non-matching LSR control, and the efficiency of the LSR was determined by the difference in mCherry percentage between the non-matching LSR control and the experimental condition.

### Integration site mapping assay to determine human genome integration specificity

#### Computational analysis of integration site mapping sequencing assay

Snakemake workflows were constructed and used to analyze NGS data from the integration site mapping sequencing assay. First, stagger sequences added to primers during library preparation were removed using custom python scripts. Next, fastp was used to trim Nextera adapters from reads and to remove reads with low PHRED scores. Next, reads were aligned to both the human genome (GRCh38) and a donor plasmid sequence containing the LSR-specific attD sequence in single-end mode using BWA MEM (H. Li and Durbin 2009). Next, reads were analyzed individually using custom python scripts to identify 1) if they aligned to the donor plasmid, human genome, or both, 2) whether or not the reads began at the predicted primer, and 3) whether or not the pre-integration attachment site was intact. Reads were then filtered to only include those reads that mapped to both the donor plasmid and the human genome, those that began at the primer site, and those that did not have an intact attD sequence (if this could be determined from the length of a particular read). This filtered read set was then aligned in paired-end mode to the human genome using default settings in BWA MEM. Alignments with a mapping quality score less than 30 were removed, along with supplementary alignments and paired read alignments with an insert size longer than 1500 bp. The samtools markdup tool was used to remove potential PCR duplicates and identify unique reads for downstream analysis (H. Li et al. 2009). Next, *MGEfinder* was used to extract clipped end sequences from reads aligned to the human genome and generate a consensus sequence of the clipped ends, which represent the crossover from the human genome into the integrated attD sequence (Durrant et al. 2020). Using custom python scripts, k-mers of length 9 base pairs were extracted from these consensus sequences and compared with a subsequence of the attD plasmid extending from the original primer to 25 bp after the end of the attD attachment site. If there were no shared 9-mers, the candidate was discarded. Otherwise, consensus sequences were clipped to begin at the primer site, and these consensus sequences were then aligned back to the original attD subsequence using the biopython local alignment tool (Cock et al. 2009). Two aligned portions were extracted - the full local alignment of the consensus sequence to the attD (called the “full local alignment”), and the longest subset of the alignment that included no ambiguous bases and no gaps (called the “contiguous alignment”). To filter a final set of true insertion sites, only sites with at least 80% nucleotide identity shared between the consensus sequence and the attD subsequence in either the full local alignment or the contiguous alignment were kept. Finally, only sites with a crossover point within 15 base pairs of the predicted dinucleotide core were kept.

This approach could precisely predict integration sites, but errors in sequencing reads led to some variability in this prediction. To account for this, integration sites were combined into integration “loci” by merging all sites that were within 500 base pairs of each other, using bedtools (Quinlan and Hall 2010). This approach would merge integration events that occurred at the same site but in opposite orientations, for example. When pooling reads across biological or technical replicates, these loci were also merged if they overlapped. When measuring the relative frequency of insertion across different loci, all uniquely aligned reads (deduplicated using samtools markdup) found within each locus were counted. These were then converted into percentages for each locus by dividing by the total number of unique reads aligned to all integration loci.

Target site motifs for different LSRs could be determined from precise predictions of dinucleotide cores for all integration sites. For each integration locus, only one integration site was chosen if there were multiple, and integration sites with more reads supporting them were prioritized. Up to 30 base pairs of human genome sequence around the predicted dinucleotide core were extracted using bedtools, choosing the forward or reverse strand depending on the orientation of the integration. All such target sites, or a subset of these target sites if desired, were then analyzed for conservation at each nucleotide position using the ggseqlogo package in R (Wagih 2017).

#### Comparison of LSR and PiggyBac transposase efficiency

1.2×10^6^ K562 cells were electroporated in 100 μL Amaxa solution (Lonza Nucleofector 2b, program T-016) with 2000 ng of a pEF-1α-PuroR-P2A-mCherry donor plasmid containing an upstream Cp36 attD site (pJT371), in combination with 3000 ng of Cp36 expression vector. Cells were grown for 10 days, then analyzed using flow cytometry for mCherry fluorescence (BioRad ZE5) with analysis using CytoFlow (https://github.com/cytoflow/cytoflow).

#### Assessment of Cp36 directionality via redosing

To generate stable mCherry-integrated cells using Cp36, 1.2×10^6^ K562 cells were electroporated in 100 μL Amaxa solution (Lonza Nucleofector 2b, program T-016) with 2000 ng of the same Cp36 PuroR-P2A-mCherry donor, in combination with 3000 ng of Cp36 expression vector. After three weeks of growth to allow the donor plasmid to dilute, cells with integrants were selected to purity using 1 μg/mL puromycin over 7 days and confirmed using flow cytometry for mCherry fluorescence (Attune NxT). To assess the efficiency of integrating a second donor sequence, we generated a second fluorescent donor construct (pJT396) by replacing mCherry in pJT371 with mTagBFP2. We then electroporated 4.0×10^5^ of wildtype or the stably integrated mCherry K562 cell lines in 100 μL Amaxa solution (Lonza Nucleofector 2b, program T-016) with pJT396 in combination with an equimolar amount of either pUC19 or a Cp36 expression vector, totalling approximately 4 μg of DNA. The frequency of doubly integrated cells was assessed using flow cytometry for mCherry and mTagBFP2 fluorescence at 13 days post-electroporation (Attune NxT), with analysis performed in FlowJo.

#### Activity assay of synthetic enhancer reporters installed at AAVS1

To install the synthetic transcription factor rTetR-VP48 into WT K562, 1.0×10^6^ WT K562 were electroporated in 100 μL Amaxa solution (Lonza Nucleofector 2b, program T-016) with 1 μg of PiggyBac expression vector (PB200A-1, SBI) and 1 μg of pMMH4, an ITR-flanked plasmid harboring the EF-1α core promoter driving rTetR-VP48-T2A-hygromycin resistance gene and a separate Tet responsive promoter (TRE3G) driving an mCherry gene. Integrants were selected to purity using 200 μg/mL hygromycin (Thermo Fisher) over 7 days. Enhancer reporter donor constructs flanked by AAVS1 homology arms (pMMH23,24,26) were subsequently integrated into the AAVS1 locus of cells expressing rTetR-VP48 using TALEN-mediated homology-directed repair as follows: 1.0×10^6^ K562 cells were electroporated in Amaxa solution (Lonza Nucleofector 2b, setting T0-16) with 1000 ng of reporter and 500 ng of each TALEN-L (Addgene #35431) and TALEN-R (Addgene #35432) plasmid (targeting upstream and downstream the intended DNA cleavage site, respectively). In the pooled reporter assay, a small library of Tet responsive elements were ordered as an oligo pool (opJS2, IDT), assembled into the reporter plasmid, miniprepped, and electroporated as a pool. Two days after electroporation, the cells were treated with 1 ng/mL puromycin antibiotic for 7 days to select for a population with reporter donor integrated into AAVS1. Reporter expression was measured by flow cytometry (BioRad ZE5) after 2 days of 1000 ng/mL doxycycline induction (Fisher Scientific).

#### Activity assay of synthetic enhancer reporters installed at a landing pad

To install the synthetic transcription factor rTetR-VP48 into landing pad cells, 1.0×10^6^ clonal Kp03 landing pad cells were electroporated in 100 μL Amaxa solution (Lonza Nucleofector 2b, program T-016) with 1 μg of PiggyBac expression vector (PB200A-1, SBI) and 1 μg of pMMH4 and selected to purity using 200 μg/mL hygromycin (Thermo Fisher) over 7 days. To install enhancer reporter plasmids at the landing pad, 1.0×10^6^ K562 cells harboring a monoclonal Kp03 landing pad and multiclonal rTetR-VP48 expression construct were electroporated in 100 μL Amaxa solution (Lonza Nucleofector 2b, program T-016) with 1000 ng of reporter donor plasmid (pMMH56,59,59). In the pooled reporter assay, 200 ng of each of five reporter constructs (pMMH55-59) were combined and electroporated together. As a negative control, cells were electroporated with 1000 ng of reporter donor with no AttP site upstream of the promoterless puro resistance gene. Three days after electroporation, the cells were treated with 1 ng/mL puromycin antibiotic for 7 days to select for a population with reporter donor correctly integrated into the landing pad. All negative control cells died during selection. Reporter expression was measured at the end of selection by flow cytometry (BioRad ZE5) after 2 days of 1000 ng/mL doxycycline induction (Fisher Scientific).

#### Magnetically separating cells based on reporter expression level

The reporter included a synthetic surface marker, consisting of the human IgG1 Fc region linked to an Igk leader and PDGFRb transmembrane domain, to enable magnetic separation of OFF from ON cells, which we previously used to study transcriptional effector domains (Tycko et al. 2020) and here adapted to study enhancers. Prior to magnetic separation, the cells were cultured between 2×10^5^ cells/mL and 1×10^6^ cells/mL for 2 weeks after selection. After 2 days of 1000 ng/mL doxycycline induction, 1×10^7^ cells were spun down at 300×g for 5 minutes and media was aspirated. Cells were then resuspended in the same volume of PBS (Gibco) and the spin down and aspiration was repeated to wash the cells and remove any IgG from serum. Dynabeads M-280 Protein G (ThermoFisher, 10003D) were resuspended by vortexing for 30 s. 50 mL of blocking buffer was prepared by adding 1 g of biotin-free BSA (Sigma Aldrich) and 200 μL of 0.5 M pH 8.0 EDTA (ThermoFisher, 15575020) into DPBS (Gibco), vacuum filtering with 0.22 μm filter (Millipore), and then kept on ice. 50 μL of beads was prepared for every 1×10^7^ cells, by adding 1 mL of buffer per 200 μL of beads, vortexing for 5 s, placing on a magnetic tube rack (Eppendorf), waiting one minute, removing supernatant, and finally removing the beads from the magnet and resuspending in 100 - 600 μL of blocking buffer per initial 50 μL of beads. Beads were added to cells at 1×10^7^ cells per 25 μL of resuspended beads, and then incubated at room temperature while rocking for 30 minutes. We used non-stick Ambion 1.5 mL tubes and a small magnetic rack. After incubation, the bead and cell mixture were placed on the magnetic rack for > 2 minutes. The unbound supernatant was transferred to a new tube, placed on the magnet again for > 2 minutes to remove any remaining beads, and then the supernatant was transferred to a new tube. For the LSR PRA, the same magnetic separation procedure was performed two more times (for a total of 3 times) on this supernatant to remove cells with activated reporters from the unbound population. Only the final unbound population was saved for further analysis by flow cytometry and library preparation. The beads from the first round of magnetic separation were resuspended in the same volume of blocking buffer, magnetically separated again, the supernatant was discarded. Resuspension, magnetic separation, and discarding the supernatant was repeated, and the tube with the beads was kept as the bound fraction. The bound fraction was resuspended in blocking buffer or PBS to dilute the cells (the unbound fraction is already dilute). Flow cytometry (BioRad ZE5) was performed using a small portion of each fraction to estimate the number of cells in each fraction and to confirm separation based on reporter levels. Finally, the samples were spun down and the pellets were frozen at 20°C until genomic DNA extraction.

#### Library preparation and sequencing of magnetically separated reporter cell pool

Genomic DNA was extracted using Monarch Genomic DNA Purification Kit (NEB) according to manufacturer instructions. After cell lysis, magnetic separation was performed on the bound population to remove beads. No more than 5×10^6^ cells were loaded onto a single column and eluted with H_2_O to avoid subsequent PCR inhibition. Libraries were assembled using 3 PCRs: PCR1 amplifies enhancer elements off the genome, PCR2 extends these amplicons with TruSeq R1/R2 handle sequences, and PCR3 extends these amplicons to add sample barcodes and p5/p7 sequences. PCR1 reactions contained 20 μL of purified gDNA, 2.5 μL of each 10 μM primer (cTF98 and cTF109), and 25 μL of Q5 2X Master Mix (NEB) and was amplified with the following thermocycling conditions: 3 minutes at 98°C, then 23X cycles of 10 seconds at 98°C, 30 seconds at 66°C, and 1 minute at 72°C, and then a final extension step of 72°C for 5 minutes. The PCR product was purified using 45 uL SPRI beads (Beckman Coulter) (0.9X of PCR volume) according to manufacturer instructions and eluted in 21 uL of nuclease free H_2_O. PCR 2 reactions were assembled with 1 μL of purified PCR 1 product, 1 μL of each 10 μM primer (oBD55 and oBD68), 10 μL of Q5 2X Master Mix, and 7 μL of nuclease-free H_2_O and amplified using the following thermocycling conditions: 30 seconds at 98°C, then 3-7X cycles of 10 seconds at 98°C, 30 seconds at 68°C, 20 seconds at 72°C, and then a final step of 72°C for 5 minutes. The PCR 2 product was purified using 18 uL SPRI beads (0.9X of PCR volume) according to manufacturer instructions and eluted in 21 uL of nuclease free H_2_O. PCR 3 reactions contained 1 μL of purified PCR 2 product, 1 μL of each 10 μM primer (oBD19-26), 10 μL of Q5 2X Master Mix, and 7 μL of nuclease-free H_2_O. The same thermocycling and purification protocol from PCR2 was performed. Purified PCR3 products were confirmed to be the correct size using a D1000 TapeStation (Agilent) and quantified with a Qubit HS kit. Samples were pooled with PhiX (Illumina) to ensure appropriate library complexity and sequenced on an Illumina Miseq with a Nano kit with 4-8 indexing cycles and 150 cycle paired-end reads.

#### Analysis of PRA sequencing data

Sequencing reads were demultiplexed using bcl2fastq (Illumina). The HT-recruit-Analyze processing pipeline was used to generate a Bowtie reference and modified to align paired-end reads with 0 mismatch allowance (https://github.com/bintulab/HT-recruit-Analyze). Count matrices for the bound and unbound samples were then used to calculate log_2_(ON:OFF) for each enhancer, normalizing for read depth across bound and unbound samples.

## External datasets

Bacterial isolate genome sequences were retrieved from GenBank and NCBI RefSeq.

## Code availability

The code for analyses performed in this paper will be available on GitHub.

## Data availability

Illumina sequencing datasets will be available on NCBI Sequence Read Archive, Bioproject PRJNA778877.

## Material availability

Requests for resources and reagents will be fulfilled by Dr. Patrick Hsu.

## Acknowledgements

We thank Aravind Natarajan, William J. Greenleaf, and the SPARK at Stanford advisors for helpful discussions. The research was supported by the Rose Hill Innovators Program at UC Berkeley and Stanford Maternal and Child Health Research Institute through Stanford’s SPARK Translational Research Program. M.G.D. is supported by the NSF GRFP and the Rose Hill Innovators program. J.T. is supported by the NIDDK F99/K00 fellowship of the NIH (F99DK126120). A.F. is supported by the NSF GRFP (2019284848). M.C.B. is supported by a grant from Stanford ChEM-H and an NIH Director’s New Innovator Award (1DP2HD084a06901). A.S.B. is supported by grants from NIH R01AI148623 and R01AI143757. This work was also supported by a grant from NIH/ENCODE 5UM1HG009436-02 (M.C.B.), NIH/NIGMS R35M128947 (L.B.). P.D.H. is supported by grants from NIH/OD DP5OD021369, NIH/NIGMS R01GM131073, DARPA, the Rainwater Foundation, the Curci Foundation, and Emergent Ventures.

## Author Contributions

M.G.D. performed bioinformatic analyses. M.G.D., J.T., A.F., M.H., and P.D.H. designed experiments. A.F., J.T., M.G.D., M.H., S.S.C., J.S., N.T.P., P.P.D, and P.D.H. performed experiments. A.F., J.T., M.G.D., M.H., J.S., and P.P.D. analyzed experimental data. M.G.D., J.T., A.F., and P.D.H. wrote the manuscript with contributions from M.C.B., L.B., A.S.B., and all other authors. M.C.B., L.B., A.S.B., and P.D.H. supervised the research.

## References

Adams, Vicki, Isabelle S. Lucet, Dena Lyras, and Julian I. Rood. 2004. “DNA Binding Properties of TnpX Indicate That Different Synapses Are Formed in the Excision and Integration of the Tn4451 Family.” Molecular Microbiology 53 (4): 1195–1207.

Almeida, Alexandre, Stephen Nayfach, Miguel Boland, Francesco Strozzi, Martin Beracochea, Zhou Jason Shi, Katherine S. Pollard, et al. 2021. “A Unified Catalog of 204,938 Reference Genomes from the Human Gut Microbiome.” Nature Biotechnology 39 (1): 105–14.

Anzalone, Andrew, Xin Gao, Christopher Podracky, Andrew Nelson, Luke Koblan, Aditya Raguram, Jonathan Levy, Jaron Mercer, and David Liu. 2021. “Programmable Large DNA Deletion, Replacement, Integration, and Inversion with Twin Prime Editing and Site-Specific Recombinases.” bioRxiv. https://doi.org/10.1101/2021.11.01.466790.

Asnicar, Francesco, Andrew Maltez Thomas, Francesco Beghini, Claudia Mengoni, Serena Manara, Paolo Manghi, Qiyun Zhu, et al. 2020. “Precise Phylogenetic Analysis of Microbial Isolates and Genomes from Metagenomes Using PhyloPhlAn 3.0.” Nature Communications 11 (1): 2500.

Calos, Michele P. 2006. “The phiC31 Integrase System for Gene Therapy.” Current Gene Therapy 6 (6): 633–45.

Cao, Jicong, Eva Maria Novoa, Zhizhuo Zhang, William C. W. Chen, Dianbo Liu, Gigi C. G. Choi, Alan S. L. Wong, Claudia Wehrspaun, Manolis Kellis, and Timothy K. Lu. 2021. “High-Throughput 5’ UTR Engineering for Enhanced Protein Production in Non-Viral Gene Therapies.” Nature Communications 12 (1): 4138.

Chalberg, Thomas W., Joylette L. Portlock, Eric C. Olivares, Bhaskar Thyagarajan, Patrick J. Kirby, Robert T. Hillman, Juergen Hoelters, and Michele P. Calos. 2006. “Integration Specificity of Phage ϕC31 Integrase in the Human Genome.” Journal of Molecular Biology 357 (1): 28–48.

Chow, Ke-Huan K., Mark W. Budde, Alejandro A. Granados, Maria Cabrera, Shinae Yoon, Soomin Cho, Ting-Hao Huang, et al. 2021. “Imaging Cell Lineage with a Synthetic Digital Recording System.” Science 372 (6538). https://doi.org/10.1126/science.abb3099.

Cock, Peter J. A., Tiago Antao, Jeffrey T. Chang, Brad A. Chapman, Cymon J. Cox, Andrew Dalke, Iddo Friedberg, et al. 2009. “Biopython: Freely Available Python Tools for Computational Molecular Biology and Bioinformatics.” Bioinformatics 25 (11): 1422–23.

Danner, Eric. n.d. “Tn5 Library Prep for Deep Sequencing Loci of CRISPR/Cas9 Edited Cells. Single Gene Specific Primer Amplification (UDiTaS Protocol with Alterations) v1.” Protocols.io. https://doi.org/10.17504/protocols.io.7k2hkye.

De Ravin, Suk See, Julie Brault, Ronald J. Meis, Siyuan Liu, Linhong Li, Mara Pavel-Dinu, Cicera R. Lazzarotto, et al. 2021. “Enhanced Homology-Directed Repair for Highly Efficient Gene Editing in Hematopoietic Stem/progenitor Cells.” Blood 137 (19): 2598–2608.

Duportet, Xavier, Liliana Wroblewska, Patrick Guye, Yinqing Li, Justin Eyquem, Julianne Rieders, Tharathorn Rimchala, Gregory Batt, and Ron Weiss. 2014. “A Platform for Rapid Prototyping of Synthetic Gene Networks in Mammalian Cells.” Nucleic Acids Research 42 (21): 13440–51.

Durrant, Matthew G., Michelle M. Li, Benjamin A. Siranosian, Stephen B. Montgomery, and Ami S. Bhatt. 2020. “A Bioinformatic Analysis of Integrative Mobile Genetic Elements Highlights Their Role in Bacterial Adaptation.” Cell Host & Microbe 28 (5): 767.

Esvelt, Kevin M., and Harris H. Wang. 2013. “Genome-Scale Engineering for Systems and Synthetic Biology.” Molecular Systems Biology 9: 641.

Farruggio, Alfonso P., and Michele P. Calos. 2014. “Serine Integrase Chimeras with Activity in E. Coli and HeLa Cells.” Biology Open 3 (10): 895–903.

Faure, Guilhem, Sergey A. Shmakov, Winston X. Yan, David R. Cheng, David A. Scott, Joseph E. Peters, Kira S. Makarova, and Eugene V. Koonin. 2019. “CRISPR-Cas in Mobile Genetic Elements: Counter-Defence and beyond.” Nature Reviews. Microbiology 17 (8): 513–25.

Fogg, Paul C. M., Ellen Younger, Booshini D. Fernando, Thanafez Khaleel, W. Marshall Stark, and Margaret C. M. Smith. 2018. “Recombination Directionality Factor gp3 Binds ϕC31 Integrase via the Zinc Domain, Potentially Affecting the Trajectory of the Coiled-Coil Motif.” Nucleic Acids Research 46 (3): 1308–20.

Ghosh, Pallavi, Amy I. Kim, and Graham F. Hatfull. 2003. “The Orientation of Mycobacteriophage Bxb1 Integration Is Solely Dependent on the Central Dinucleotide of attP and attB.” Molecular Cell 12 (5): 1101–11.

Giannoukos, Georgia, Dawn M. Ciulla, Eugenio Marco, Hayat S. Abdulkerim, Luis A. Barrera, Anne Bothmer, Vidya Dhanapal, et al. 2018. “UDiTaS™, a Genome Editing Detection Method for Indels and Genome Rearrangements.” BMC Genomics 19 (1): 212.

Groth, A. C., E. C. Olivares, B. Thyagarajan, and M. P. Calos. 2000. “A Phage Integrase Directs Efficient Site-Specific Integration in Human Cells.” Proceedings of the National Academy of Sciences of the United States of America 97 (11): 5995–6000.

Guha, Tuhin K., and Michele P. Calos. 2020. “Nucleofection of phiC31 Integrase Protein Mediates Sequence-Specific Genomic Integration in Human Cells.” Journal of Molecular Biology 432 (13): 3950–55.

Haapaniemi, Emma, Sandeep Botla, Jenna Persson, Bernhard Schmierer, and Jussi Taipale. 2018. “CRISPR-Cas9 Genome Editing Induces a p53-Mediated DNA Damage Response.” Nature Medicine 24 (7): 927–30.

Haberle, Vanja, Cosmas D. Arnold, Michaela Pagani, Martina Rath, Katharina Schernhuber, and Alexander Stark. 2019. “Transcriptional Cofactors Display Specificity for Distinct Types of Core Promoters.” Nature 570 (7759): 122–26.

Hauser, Maria, Martin Steinegger, and Johannes Söding. 2016. “MMseqs Software Suite for Fast and Deep Clustering and Searching of Large Protein Sequence Sets.” Bioinformatics 32 (9): 1323–30.

Inniss, Mara C., Kalpanie Bandara, Barbara Jusiak, Timothy K. Lu, Ron Weiss, Liliana Wroblewska, and Lin Zhang. 2017. “A Novel Bxb1 Integrase RMCE System for High Fidelity Site-Specific Integration of mAb Expression Cassette in CHO Cells.” Biotechnology and Bioengineering 114 (8): 1837–46.

Ioannidi, Eleonora I., Matthew T. N. Yarnall, Cian Schmitt-Ulms, Rohan N. Krajeski, Justin Lim, Lukas Villiger, Wenyuan Zhou, et al. 2021. “Drag-and-Drop Genome Insertion without DNA Cleavage with CRISPR-Directed Integrases.” bioRxiv. https://doi.org/10.1101/2021.11.01.466786.

Jusiak, Barbara, Kalpana Jagtap, Leonid Gaidukov, Xavier Duportet, Kalpanie Bandara, Jianlin Chu, Lin Zhang, Ron Weiss, and Timothy K. Lu. 2019. “Comparison of Integrases Identifies Bxb1-GA Mutant as the Most Efficient Site-Specific Integrase System in Mammalian Cells.” ACS Synthetic Biology 8 (1): 16–24.

Karow, Marisa, Christopher L. Chavez, Alfonso P. Farruggio, Jonathan M. Geisinger, Annahita Keravala, W. Edward Jung, Feng Lan, Joseph C. Wu, Yanru Chen-Tsai, and Michele P. Calos. 2011. “Site-Specific Recombinase Strategy to Create Induced Pluripotent Stem Cells Efficiently with Plasmid DNA.” Stem Cells 29 (11): 1696–1704.

Katoh, Kazutaka, and Daron M. Standley. 2013. “MAFFT Multiple Sequence Alignment Software Version 7: Improvements in Performance and Usability.” Molecular Biology and Evolution 30 (4): 772–80.

Keravala, Annahita, Joylette L. Portlock, Joan A. Nash, David G. Vitrant, Paul D. Robbins, and Michele P. Calos. 2006. “PhiC31 Integrase Mediates Integration in Cultured Synovial Cells and Enhances Gene Expression in Rabbit Joints.” The Journal of Gene Medicine 8 (8): 1008–17.

Khaleel, Thanafez, Ellen Younger, Andrew R. McEwan, Anpu S. Varghese, and Margaret C. M. Smith. 2011. “A Phage Protein That Binds φC31 Integrase to Switch Its Directionality.” Molecular Microbiology 80 (6): 1450–63.

King, Dana M., Clarice Kit Yee Hong, James L. Shepherdson, David M. Granas, Brett B. Maricque, and Barak A. Cohen. 2020. “Synthetic and Genomic Regulatory Elements Reveal Aspects of Cis-Regulatory Grammar in Mouse Embryonic Stem Cells.” eLife 9 (February). https://doi.org/10.7554/eLife.41279.

Klein, Jason C., Vikram Agarwal, Fumitaka Inoue, Aidan Keith, Beth Martin, Martin Kircher, Nadav Ahituv, and Jay Shendure. 2020. “A Systematic Evaluation of the Design and Context Dependencies of Massively Parallel Reporter Assays.” Nature Methods 17 (11): 1083–91.

Kosicki, Michael, Kärt Tomberg, and Allan Bradley. 2018. “Repair of Double-Strand Breaks Induced by CRISPR-Cas9 Leads to Large Deletions and Complex Rearrangements.” Nature Biotechnology 36 (8): 765–71.

Kung, Stephanie H., Adam C. Retchless, Jessica Y. Kwan, and Rodrigo P. P. Almeida. 2013. “Effects of DNA Size on Transformation and Recombination Efficiencies in Xylella Fastidiosa.” Applied and Environmental Microbiology 79 (5): 1712–17.

Lee, Jae Seong, Lise Marie Grav, Lasse Ebdrup Pedersen, Gyun Min Lee, and Helene Faustrup Kildegaard. 2016. “Accelerated Homology-Directed Targeted Integration of Transgenes in Chinese Hamster Ovary Cells via CRISPR/Cas9 and Fluorescent Enrichment.” Biotechnology and Bioengineering 113 (11): 2518–23.

Li, Heng, and Richard Durbin. 2009. “Fast and Accurate Short Read Alignment with Burrows–Wheeler Transform.” Bioinformatics 25 (14): 1754–60.

Li, Heng, Bob Handsaker, Alec Wysoker, Tim Fennell, Jue Ruan, Nils Homer, Gabor Marth, Goncalo Abecasis, Richard Durbin, and 1000 Genome Project Data Processing Subgroup. 2009. “The Sequence Alignment/Map Format and SAMtools.” Bioinformatics 25 (16): 2078–79.

Li, Xianghong, Erin R. Burnight, Ashley L. Cooney, Nirav Malani, Troy Brady, Jeffry D. Sander, Janice Staber, et al. 2013. “piggyBac Transposase Tools for Genome Engineering.” Proceedings of the National Academy of Sciences of the United States of America 110 (25): E2279–87.

Maricque, Brett B., Hemangi G. Chaudhari, and Barak A. Cohen. 2018. “A Massively Parallel Reporter Assay Dissects the Influence of Chromatin Structure on Cis-Regulatory Activity.” Nature Biotechnology, November. https://doi.org/10.1038/nbt.4285.

Matreyek, Kenneth A., Jason J. Stephany, Melissa A. Chiasson, Nicholas Hasle, and Douglas M. Fowler. 2020. “An Improved Platform for Functional Assessment of Large Protein Libraries in Mammalian Cells.” Nucleic Acids Research 48 (1): e1.

McEwan, Andrew R., Paul A. Rowley, and Margaret C. M. Smith. 2009. “DNA Binding and Synapsis by the Large C-Terminal Domain of phiC31 Integrase.” Nucleic Acids Research 37 (14): 4764–73.

Merrick, Christine A., Jia Zhao, and Susan J. Rosser. 2018. “Serine Integrases: Advancing Synthetic Biology.” ACS Synthetic Biology 7 (2): 299–310.

Perez, Christophe, Valérie Guyot, Jean-Pierre Cabaniols, Agnès Gouble, Beatrice Micheaux, Julie Smith, Sophie Leduc, Frédéric Pâques, and Philippe Duchateau. 2005. “Factors Affecting Double-Strand Break-Induced Homologous Recombination in Mammalian Cells.” BioTechniques 39 (1): 109–15.

Ptáčková, Pavlína, Jan Musil, Martin Štach, Petr Lesný, Šárka Němečková, Vlastimil Král, Milan Fábry, and Pavel Otáhal. 2018. “A New Approach to CAR T-Cell Gene Engineering and Cultivation Using piggyBac Transposon in the Presence of IL-4, IL-7 and IL-21.” Cytotherapy 20 (4): 507–20.

Quinlan, Aaron R., and Ira M. Hall. 2010. “BEDTools: A Flexible Suite of Utilities for Comparing Genomic Features.” Bioinformatics 26 (6): 841–42.

Rabani, Michal, Lindsey Pieper, Guo-Liang Chew, and Alexander F. Schier. 2017. “A Massively Parallel Reporter Assay of 3’ UTR Sequences Identifies In Vivo Rules for mRNA Degradation.” Molecular Cell 68 (6): 1083–94.e5.

Salmond, George P. C., and Peter C. Fineran. 2015. “A Century of the Phage: Past, Present and Future.” Nature Reviews. Microbiology 13 (12): 777–86.

Sandoval-Villegas, Nicolás, Wasifa Nurieva, Maximilian Amberger, and Zoltán Ivics. 2021. “Contemporary Transposon Tools: A Review and Guide through Mechanisms and Applications of Sleeping Beauty, piggyBac and Tol2 for Genome Engineering.” International Journal of Molecular Sciences 22 (10). https://doi.org/10.3390/ijms22105084.

Sclimenti, C. R., B. Thyagarajan, and M. P. Calos. 2001. “Directed Evolution of a Recombinase for Improved Genomic Integration at a Native Human Sequence.” Nucleic Acids Research 29 (24): 5044–51.

Shendure, Jay, Shankar Balasubramanian, George M. Church, Walter Gilbert, Jane Rogers, Jeffery A. Schloss, and Robert H. Waterston. 2017. “DNA Sequencing at 40: Past, Present and Future.” Nature 550 (7676): 345–53.

Sivalingam, Jaichandran, Shruti Krishnan, Wai Har Ng, Sze Sing Lee, Toan Thang Phan, and Oi Lian Kon. 2010. “Biosafety Assessment of Site-Directed Transgene Integration in Human Umbilical Cord-Lining Cells.” Molecular Therapy: The Journal of the American Society of Gene Therapy 18 (7): 1346–56.

Sivalingam, J., S. Krishnan, W. H. Ng, S. S. Lee, and T. T. Phan. 2010. “Biosafety Assessment of Site-Directed Transgene Integration in Human Umbilical Cord–lining Cells.” Molecular Therapy: The Journal of the American Society of Gene Therapy. https://www.sciencedirect.com/science/article/pii/S1525001616310802.

Smith, Margaret C. M. 2015. “Phage-Encoded Serine Integrases and Other Large Serine Recombinases.” Mobile DNA III, 253–72.

Smith, Margaret C. M., and Helena M. Thorpe. 2002. “Diversity in the Serine Recombinases.” Molecular Microbiology 44 (2): 299–307.

Steinegger, Martin, and Johannes Söding. 2017. “MMseqs2 Enables Sensitive Protein Sequence Searching for the Analysis of Massive Data Sets.” Nature Biotechnology 35 (11): 1026–28.

Thyagarajan, Bhaskar, Ying Liu, Soojung Shin, Uma Lakshmipathy, Kelly Scheyhing, Haipeng Xue, Catharina Ellerström, et al. 2008. “Creation of Engineered Human Embryonic Stem Cell Lines Using phiC31 Integrase.” Stem Cells 26 (1): 119–26.

Tycko, Josh, Nicole DelRosso, Gaelen T. Hess Aradhana, Abhimanyu Banerjee, Aditya Mukund, Mike V. Van, et al. 2020. “High-Throughput Discovery and Characterization of Human Transcriptional Effectors.” Cell 183 (7): 2020–35.e16.

Vaidyanathan, Sriram, Ron Baik, Lu Chen, Dawn T. Bravo, Carlos J. Suarez, Shayda M. Abazari, Ameen A. Salahudeen, et al. 2021. “Targeted Replacement of Full-Length CFTR in Human Airway Stem Cells by CRISPR-Cas9 for Pan-Mutation Correction in the Endogenous Locus.” Molecular Therapy: The Journal of the American Society of Gene Therapy, March. https://doi.org/10.1016/j.ymthe.2021.03.023.

Wagih, Omar. 2017. “Ggseqlogo: A Versatile R Package for Drawing Sequence Logos.” Bioinformatics 33 (22): 3645–47.

Wang, H., and P. Mullany. 2000. “The Large Resolvase TndX Is Required and Sufficient for Integration and Excision of Derivatives of the Novel Conjugative Transposon Tn5397.” Journal of Bacteriology 182 (23): 6577–83.

Wang Hongmei, Smith Margaret C.M., and Mullany Peter. 2006. “The Conjugative Transposon Tn5397 Has a Strong Preference for Integration into Its Clostridium Difficile Target Site.” Journal of Bacteriology 188 (13): 4871–78.

Wang, Wei, Chengyi Lin, Dong Lu, Zeming Ning, Tony Cox, David Melvin, Xiaozhong Wang, Allan Bradley, and Pentao Liu. 2008. “Chromosomal Transposition of PiggyBac in Mouse Embryonic Stem Cells.” Proceedings of the National Academy of Sciences of the United States of America 105 (27): 9290–95.

Wang, Xianwei, Biao Tang, Yu Ye, Yayi Mao, Xiaolai Lei, Guoping Zhao, and Xiaoming Ding. 2017. “Bxb1 Integrase Serves as a Highly Efficient DNA Recombinase in Rapid Metabolite Pathway Assembly.” Acta Biochimica et Biophysica Sinica 49 (1): 44–50.

Weingarten-Gabbay, Shira, Ronit Nir, Shai Lubliner, Eilon Sharon, Yael Kalma, Adina Weinberger, and Eran Segal. 2019. “Systematic Interrogation of Human Promoters.” Genome Research 29 (2): 171–83.

Wilson, Matthew H., Craig J. Coates, and Alfred L. George Jr. 2007. “PiggyBac Transposon-Mediated Gene Transfer in Human Cells.” Molecular Therapy: The Journal of the American Society of Gene Therapy 15 (1): 139–45.

Yang, Lei, Alec A. K. Nielsen, Jesus Fernandez-Rodriguez, Conor J. McClune, Michael T. Laub, Timothy K. Lu, and Christopher A. Voigt. 2014. “Permanent Genetic Memory With> 1-Byte Capacity.” Nature Methods 11 (12): 1261–66.

Yusa, Kosuke. 2015. “piggyBac Transposony.” In Mobile DNA III, 873–90. Washington, DC, USA: ASM Press.

Yusa, Kosuke, Liqin Zhou, Meng Amy Li, Allan Bradley, and Nancy L. Craig. 2011. “A Hyperactive piggyBac Transposase for Mammalian Applications.” Proceedings of the National Academy of Sciences of the United States of America 108 (4): 1531–36.

